# Designing Efficient Enzymes: Eight Predicted Mutations Convert a Hydroxynitrile Lyase into an Efficient Esterase

**DOI:** 10.1101/2023.08.23.554512

**Authors:** Guillem Casadevall, Colin Pierce, Bo Guan, Javier Iglesias-Fernandez, Huey-Yee Lim, Lauren R. Greenberg, Meghan E. Walsh, Ke Shi, Wendy Gordon, Hideki Aihara, Robert L. Evans, Romas Kazlauskas, Sílvia Osuna

**Affiliations:** Institut de Química Computacional i Catàlisi and Departament de Química, Universitat de Girona, Carrer Maria Aurèlia Capmany 69, 17003 Girona, Spain; Biotechnology Institute and Department of Biochemistry, Molecular Biology and Biophysics, University of Minnesota, 1479 Gortner Avenue, Saint Paul, MN 55108 USA; ICREA, Barcelona, Spain

## Abstract

Hydroxynitrile lyase from rubber tree (*Hb*HNL) shares 45% identical amino acid residues with the homologous esterase from tobacco, SABP2, but the two enzymes catalyze different reactions. The x-ray structures reveal a serine-histidine-aspartate catalytic triad in both enzymes along with several differing amino acid residues within the active site. Previous exchange of three amino acid residues in the active site of *Hb*HNL with the corresponding amino acid residue in SABP2 (T11G-E79H-K236M) created variant HNL3, which showed low esterase activity toward p-nitrophenyl acetate. Further structure comparison reveals additional differences surrounding the active site. *Hb*HNL contains an improperly positioned oxyanion hole residue and differing solvation of the catalytic aspartate. We hypothesized that correcting these structural differences would impart good esterase activity on the corresponding HbHNL variant. To predict the amino acid substitutions needed to correct the structure, we calculated shortest path maps for both *Hb*HNL and SABP2, which reveal correlated movements of amino acids in the two enzymes. Replacing four amino acid residues (C81L-N104T-V106F-G176S) whose movements are connected to the movements of the catalytic residues yielded variant HNL7TV (stabilizing substitution H103V was also added), which showed an esterase catalytic efficiency comparable to that of SABP2. The x-ray structure of an intermediate variant, HNL6V, showed an altered solvation of the catalytic aspartate and a partially corrected oxyanion hole. This dramatic increase in catalytic efficiency demonstrates the ability of shortest path maps to predict which residues outside the active site contribute to catalytic activity.

## Introduction

One of the most important qualities of an enzyme is its ability to efficiently catalyze reactions. Nature’s enzymes are efficient with natural substrates, but many potential applications require enzymes to work with unnatural substrates or even catalyze new chemical steps. Engineering enzymes to Nature-like efficiencies remains an unsolved problem.^[1]^ Protein design often yields enzymes millions of times slower than natural enzymes.

Improving enzyme efficiency is challenging for several reasons. First, an efficient enzyme must simultaneously optimize substrate binding, transition state stabilization, and product release. Second, proteins move continuously,^[2-6]^ but catalysis requires precisely positioning the substrate and catalytic groups for reaction. It is difficult to predict how to shift the conformational landscape to favor the catalytically competent conformations. Third, residues outside the active site, not in direct contact with substrate, contribute to catalysis as shown by many directed evolution experiments, but it is difficult to predict how these distant residues impact catalysis.

The objective of this paper is to test whether correlated protein motions (shortest path maps, SPM^[1, 4]^) can predict residues outside the active site that contribute to catalysis. These SPM’s identify residues that move together during molecular dynamics simulations. We hypothesize that residues whose motion is correlated with the motions of the catalytic residues are those that contribute most strongly to efficient catalysis. Mutating these correlated positions should shift global minimum energy conformation towards catalytically competent conformations. Previously, SPM’s identified locations of beneficial substitutions previously identified by directed evolution,^[1, 4]^ and substitutions that allosterically activated tryptophan synthase B a modest 7-fold in k_cat_ and 4-fold in k_cat_/K_M_.^[7]^

The test case is to increase the efficiency of an inefficient esterase (a modified hydroxynitrile lyase, HNL3V) by computational design. The engineering uses a homologous esterase, SABP2, for comparison and focuses on transferring substitutions from the homologous esterase to the modified hydroxynitrile lyase. This approach limits the scope of the problem, but it remains challenging since the esterase and modified hydroxynitrile lyase differ by 146 residues. There are 2^146^ or approximately 10^44^ possible variants one could create by exchanging residues between the two enzymes.

Previous design of esterase activity achieved only modest catalytic efficiencies;^[8-12]^ the best k_cat_/K_M_ was 6,600 M^-1^ min^-1^.^[11]^ In contrast, the work below reports an eight-substitution variant with excellent catalytic efficiency (k_cat_/K_M_ of 120,000 M^-1^ min^-1^), which is two-fold higher than that for the target esterase. In addition, computational and x-ray structure analysis of the variants reveal the molecular basis of the improved esterase activity: improved positioning of an oxyanion-hole-stabilizing residue and altered solvation of the catalytic aspartate.

## Results

### Homologs SABP2 and *Hb*HNL catalyze different reactions with similar active sites

*Hb*HNL and SABP2 are homologous enzymes with similar catalytic triads but different catalytic activity. *Hb*HNL shares 44% sequence identity and 62% similarity over 260 positions with the modern esterase SABP2 (salicylic acid binding protein 2 from tobacco), Supplementary Fig. 1. *Hb*HNL and SABP2 contain the same Ser-His-Asp catalytic triad, but *Hb*HNL catalyzes cyanohydrin cleavage,^[13]^ while SABP2 catalyzes ester hydrolysis.^[14]^ Both enzymes belong to the ɑ/β-hydrolase fold superfamily.^[15, 16]^ *Hb*HNL and other hydroxynitrile lyases diverged from the esterases approximately 100 million years ago.^[17, 18]^

SABP2 and other serine esterases contain a catalytic triad (Ser-His-Asp) and an oxyanion hole, which consists of two main chain N-H’s that form hydrogen bonds to the substrate carbonyl oxygen, Fig. 1. In the first step of the reaction, His238 abstracts a proton from the O_γ_ of Ser81, which initiates a nucleophilic attack on the carbonyl carbon of the pNPAc substrate (Fig. 1a). The resultant oxyanion is stabilized through H-bonds with backbone N-H’s at positions 13 and 82.

**Fig. 1.**
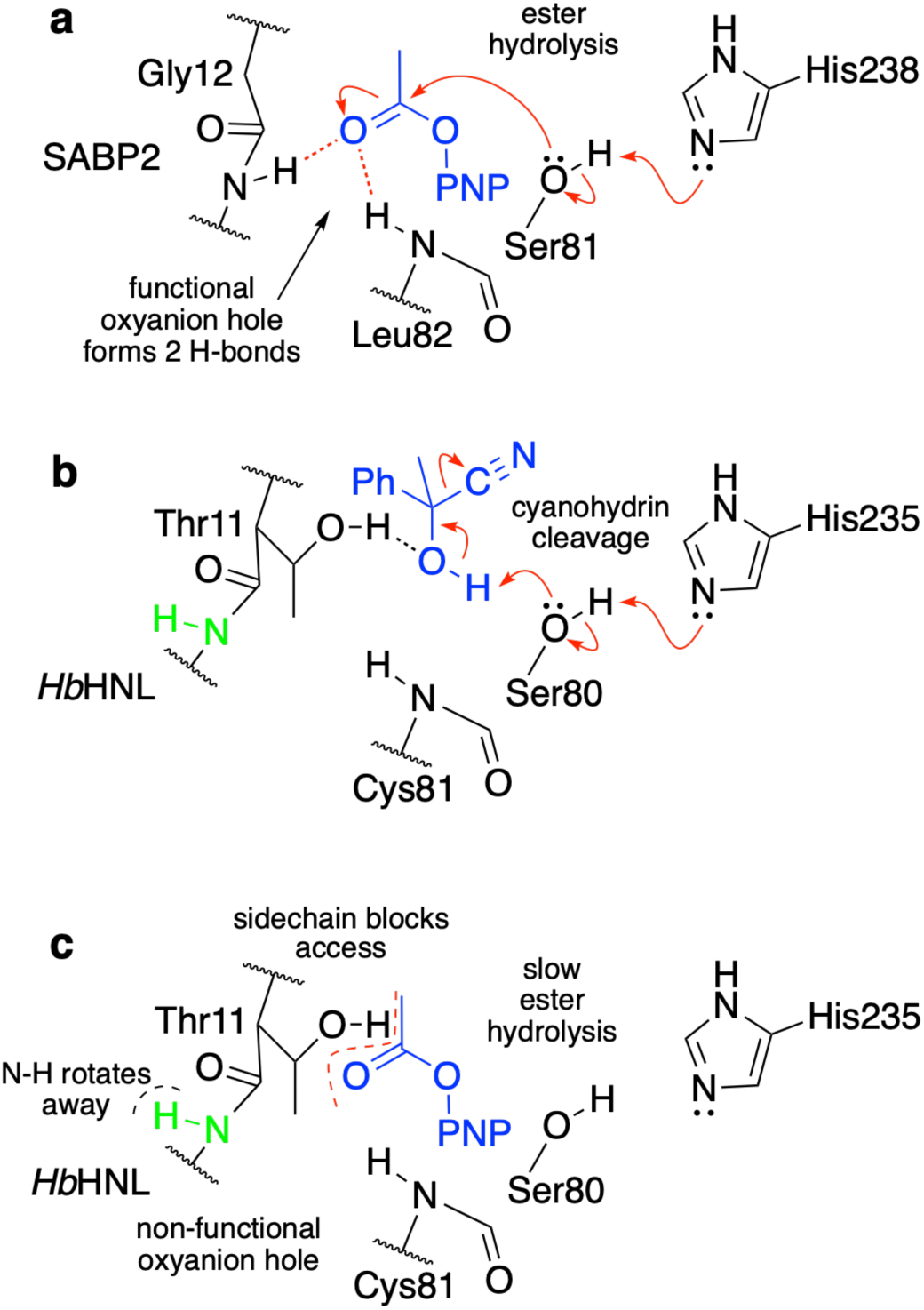
Although esterase SABP2 and hydroxynitrile lyase *Hb*HNL both contain a Ser-His-Asp catalytic triad, *Hb*HNL lacks a functional oxyanion hole preventing it from catalyzing ester hydrolysis. **a**, The carbonyl oxygen of the *p*-nitrophenyl acetate substrate (blue) accepts hydrogen bonds from two main chain N-H’s (Gly12, Leu82). These simultaneous hydrogen bonds are known as the oxyanion hole. Hydrolysis of the ester starts with a nucleophilic attack of Ser81 on the carbonyl carbon to form a tetrahedral intermediate (not shown). The oxyanion hole stabilizes the negative charge that forms on the oxygen in the tetrahedral intermediate. **b**, *Hb*HNL catalyzes cleavage of a cyanohydrin (blue) in a single step. The reaction does not involve an oxyanion intermediate, so catalysis does not require an oxyanion hole. **c**, *Hb*HNL catalyzes slow *p*-nitrophenyl acetate hydrolysis (blue). The oxyanion hole in *Hb*HNL is disrupted by the side chain of Thr11, which hinders access of the substrate to this region and by a twist in the main chain that points the N-H (green) away from the substrate. For clarity none of the diagrams show the aspartate of the catalytic triad.

*Hb*HNL catalyzes the enantioselective cleavage of mandelonitrile, an aromatic cyanohydrin (Fig. 1b).^[19, 20]^ In contrast to the multi-step ester hydrolysis catalyzed by SABP2, this lyase reaction occurs in a single step. Catalysis of mandelonitrile cleavage uses the catalytic triad for simple acid-base chemistry. The catalytic Ser deprotonates the substrate hydroxyl; subsequent elimination of cyanide yields benzaldehyde. The lyase reaction does not involve an oxyanion intermediate.

*Hb*HNL also catalyzes promiscuous hydrolysis of pNPAc, but 500-fold slower than SABP2 (k_cat_ of 0.25 min^-1^ vs. 130 min^-1^ for SABP2), Fig. 1c. One reason for the low esterase activity of *Hb*HNL is that access to the oxyanion hole is blocked by the side chain of Thr11. During lyase catalysis, the side chain hydroxyl of Thr11 can donate a hydrogen bond to the cyanohydrin hydroxyl, Fig. 1b. The corresponding residue in SABP2 is Gly12, which allows full access to the oxyanion hole. Replacement of Gly12 with threonine in SABP2 decreased the esterase activity 2000-fold^[21]^ confirming that the threonine hinders ester hydrolysis. However, replacement of Thr11 in *Hb*HNL with glycine increases the promiscuous hydrolysis of pNPAc only slightly^[19]^ suggesting that additional structural differences between the oxyanion hole in *Hb*HNL and SABP2 contribute to the low esterase activity of *Hb*HNL.

### The positioning of one oxyanion hole residue differs significantly between SABP2 and *Hb*HNL

The x-ray crystal structures of SABP2 and *Hb*HNL show similar placement for the catalytic atoms. The x-ray structure of apo SABP2 and three x-ray structures of apo *Hb*HNL were overlaid to best fit the positions of all corresponding Cɑ, Fig. 2. The comparison was restricted to structures without a ligand in the active site to avoid structure changes caused by the ligand. The comparison of multiple structures may also account for some of the motion of the atoms in solution. Four of the catalytic atoms (serine Oɣ, histidine Nε2, aspartate Oδ2, and oxyanion hole residue OX2 N) align closely between SABP2 and the three HbNHL structures with RMSD of 0.4-0.9 Å.

**Fig. 2.**
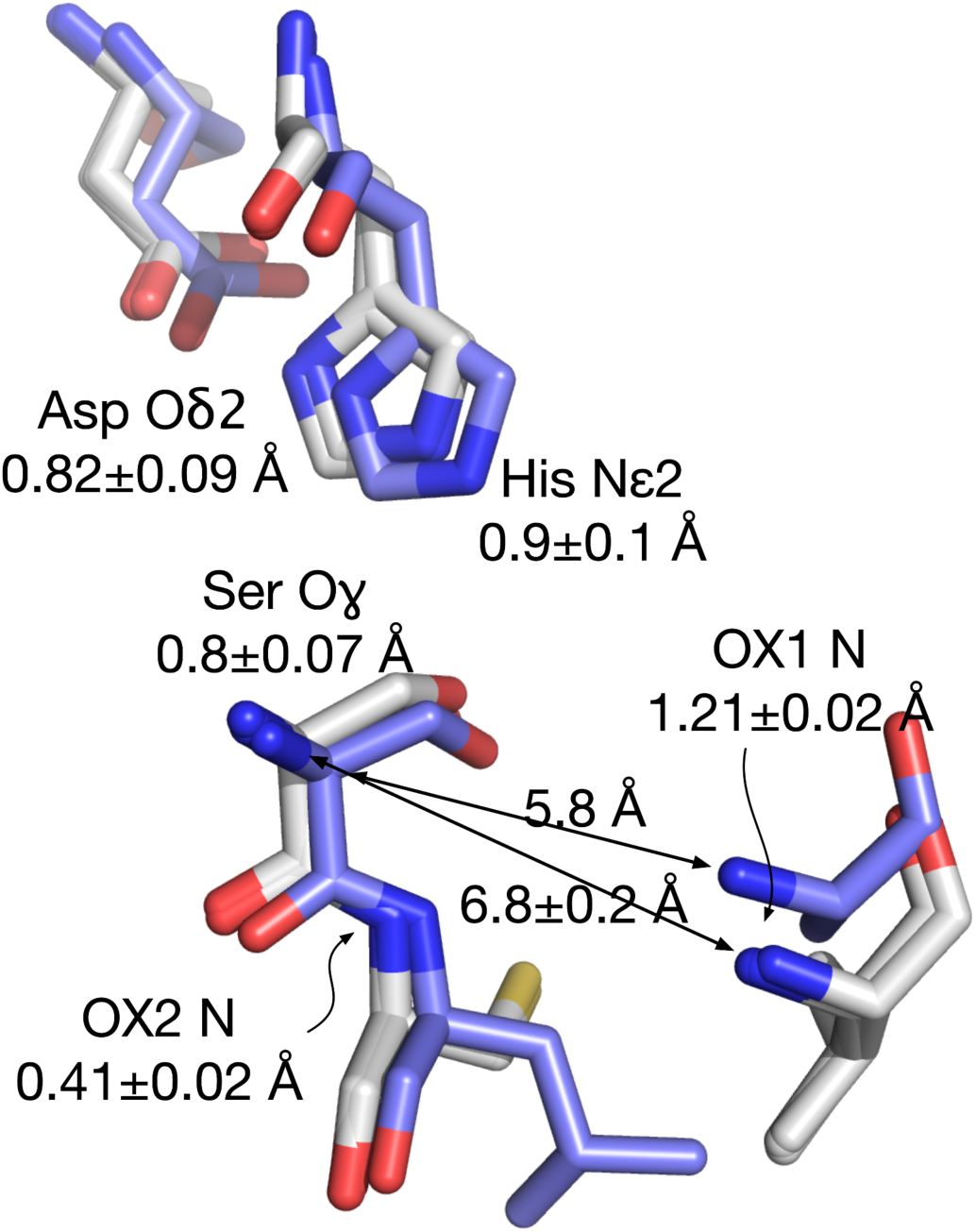
Overlay of the catalytic residues of three apo *Hb*HNL structures. (pdb id = 6yas, 3c6x, 2g4l; white carbons) onto the structure of apo SABP2 (1y7h; blue carbons). The alignment minimized the RMSD between the corresponding Cɑ atoms in the entire protein. The average deviation was 0.68 Å over 213-218 aligned Cɑ atoms out of 256. The five catalytic triad and oxyanion hole atoms deviated by slightly more than the average Cɑ deviation, an average of 0.8 ± 0.3 Å. The largest deviation was the catalytic nitrogen atom of oxyanion residue OX1 (Ala13 in SABP2, Ile12 in *Hb*HNL), which deviated by 1.21±0.02 Å. This deviation likely contributes to the poor esterase activity of *Hb*HNL. Another way to measure this difference is the serine Cɑ to OX1 N distance within each structure. For SABP2, this distance is 5.8 Å, while for the three *Hb*HNL structures, this distance is longer, 6.8 ± 0.2 Å.

The remaining catalytic atom (oxyanion hole residue OX1 N) differs significantly between SABP2 and *Hb*HNL: 1.2 Å. We hypothesized that this difference contributes to the poor promiscuous esterase activity of *Hb*HNL. The k_cat_/K_M_ value for hydrolysis of *p*-nitrophenyl acetate by *Hb*HNL is only 0.1% of the value for SABP2, Table 1. Since OX1 N is a main chain atom, fixing this deviation requires shifting the protein backbone. A disrupted oxyanion hole in *Hb*HNL is not expected to hinder lyase catalysis because it does not involve an oxyanion intermediate.

**Table 1.** Steady-state kinetics parameters for *p*-nitrophenyl acetate hydrolysis catalyzed by SABP2, *Hb*HNL and *Hb*HNL enzyme variants. See Supplementary Table 1 for a more extensive list of enzyme variants.

| Variant | $k_{\text{cat}}$ ($\text{min}^{-1}$) | $K_{\text{M}}$ (mM) | $k_{\text{cat}}/K_{\text{M}}$ ($\text{M}^{-1} \cdot \text{min}^{-1}$) | $k_{\text{cat}}$ (% relative to SABP2) | $k_{\text{cat}}/K_{\text{M}}$ (% relative to SABP2) |
| --- | --- | --- | --- | --- | --- |
| SABP2 | $134 \pm 4$ | $2.2 \pm 0.2$ | 61,000 | 100% | 100% |
| *HbHNL* | $0.25 \pm 0.02$ | $3.0 \pm 0.4$ | 83 | 0.2% | 0.1% |
| HNL3 | $0.33 \pm 0.02$ | $0.71 \pm 0.1$ | 460 | 0.2% | 0.8% |
| HNL3V | $0.32 \pm 0.02$ | $0.65 \pm 0.1$ | 490 | 0.2% | 0.9% |
| H N L 3 V<br>N 1 0 4 A -<br>G176S | $2.1 \pm 0.17$ | $0.04 \pm 0.01$ | 52,000 | 1.6% | 87% |
| HNL6V<sup>a</sup> | $0.59 \pm 0.14$ | $0.03 \pm 0.003$ | 20,000 | 0.4% | 33% |
| HNL7V | $3.9 \pm 0.5$ | $0.16 \pm 0.07$ | 25,000 | 2.9% | 40% |
| HNL7TV | $9.3 \pm 0.3$ | $0.08 \pm 0.04$ | 120,000 | 7.0% | 190% |
| HNL8V | $8.1 \pm 0.4$ | $0.35 \pm 0.06$ | 23,000 | 6.1% | 38% |

<sup>a</sup> Preliminary data for HNL6V measured at the typical temperature of 21°C. $k_{\text{cat}}$ at 29°C was $2.3 \pm 0.02 \text{ min}^{-1}$ , $K_{\text{M}}$ $0.13 \pm 0.01 \text{ mM}$

The internal distance between serine Cɑ and OX1 N is significantly longer in the *Hb*HNL structures (6.8 ± 0.2 Å) than in SABP2 (5.8 Å). This difference indicates that the ester carbonyl group, which interacts with the serine Oɣ and OX1 N in the transition state, cannot make the same interactions in *Hb*HNL and SABP2.

### Previous engineering within active site yielded only inefficient esterase activity

Three amino acid residues in the active site of *Hb*HNL are thought to have a mechanistic role in the catalysis of the lyase reaction, but hinder ester hydrolysis.^[21]^ The side chain of Thr11 helps orient the hydroxynitrile substrate, but blocks access of the ester substrate to the oxyanion hole (Fig. 1c). Lys236, oriented by Glu79, stabilizes the leaving cyanide from hydroxynitriles but hinders the loss of a hydrophobic group from an ester. Replacement of these three residues in *Hb*HNL with the corresponding residues in SABP2 (HbHNL-T11G-E79H-K236M) to create HNL3 increased the esterase catalytic efficiency (k_cat_/K_M_) of *Hb*HNL 5.6-fold from 84 to 470 *M*^-1^·min^-1^, Table 1. For experimental convenience, we created HNL3V, which contains an additional H103V substitution that stabilizes the protein. This H103V substitution did not affect the esterase activity of HNL3 and is analogous to the stabilizing H103L substitution in the homologous HNL from *Manihot esculenta*.^[22]^ We hypothesized that additional substitutions outside the active site are required to reposition the catalytic machinery of *Hb*HNL, including the oxyanion hole, to enable efficient esterase activity.

### Shortest path map identifies residues outside the active site that alter the conformational landscape

Since proteins move and flex continuously, repositioning of the catalytic atoms requires altering the conformational landscape of the enzyme. The shortest path map (SPM) methodology identified the positions that contribute most strongly to conformational dynamics in SABP2 and HNL3V and replaced those amino acids that differed between them within the SPM closest to the active site. SPM starts with a molecular dynamics simulation and identifies which residues move together and have a higher contribution to the conformational dynamics.^[1,4]^ Importantly, SPM predicts which amino acids outside the active site can alter the positioning of the catalytic groups within the active site.

The SPMs constructed for HNL3V and SABP2 revealed differences in the motions involving the catalytic residues, Fig. 3. Based on those SPM’s, two regions were identified for mutagenesis. In the first region two substitutions are expected to add a correlated motion with OX1 and in the second region three substitutions are expected to remove a correlated motion to the catalytic Asp207. The SPM of SABP2 indicates that movements of OX1 (Ala13) are directly connected to motions of residues Cys14, Gly12, Ser179, and Leu 82, Supplementary Fig. 2. The first two residues are conserved between SABP2 and HNL3V, but the second two residues differ. Positions Ser179 and Leu82 in SABP2 correspond to Gly176 and Cys81 in HNL3V. Residue Cys81 does not appear in the SPM of HNL3V indicating that its motions are not strongly correlated with the movements of any other residues including OX1 (Ile12). While the motion of Ser179 is directly correlated to OX1 (Ala13) in SABP2, the motion between OX1 (Ile12) and Gly176 in HNL3V differs because it correlates indirectly via Cys13. Therefore, SPM analysis predicted that the C81L and G176S substitutions in HNL3V would create motions directly correlated with the oxyanion hole residue Ile12 and may fix its orientation. OX1 (Ala13), which corresponds to Ile12 in HNL, is also part of the correlated motion in SABP2. We did not include an Ile12Ala substitution in the variants because the Ala13Leu substitution had no effect on esterase activity.^[23]^

**Fig. 3.**
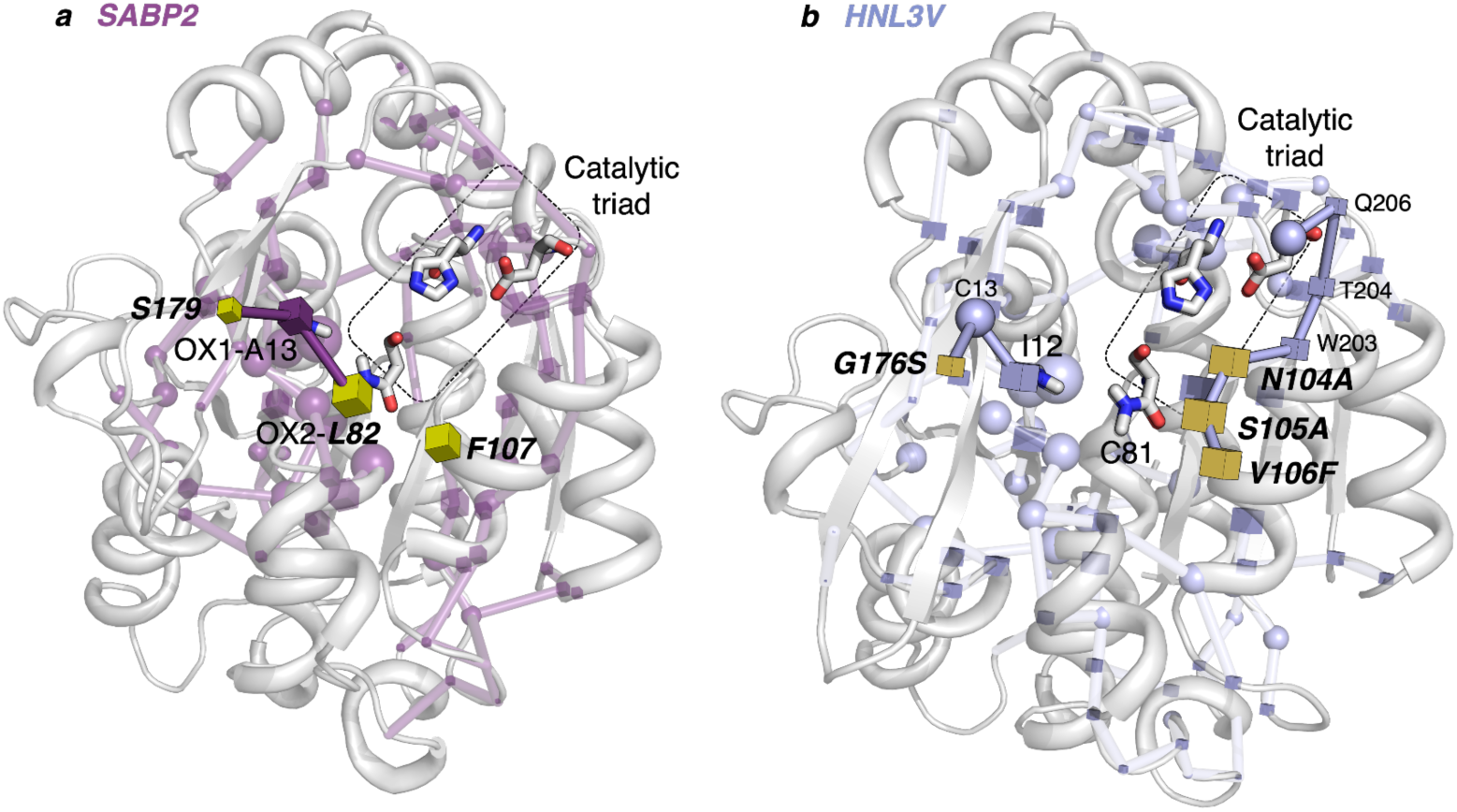
Shortest path maps of (a) SABP2 and (b) HNL3V showing five substitutions predicted to make the conformational dynamics of catalytic residues in HNL3V more like that of SABP2. The substitutions Gly176Ser and Cys81Leu add a connection to OX1 (Ile11) similar to that present in SABP2 where OX1 corresponds to Ala13. The substitutions at 104-106 in HNL3V remove a connection to the catalytic Asp207 that is present in HNL3V, but absent in SABP2. Adding these five substitutions to HNL3V created HNL8V. The spheres in the SPM indicate residues conserved between the two proteins, while cubes indicate residues that differ between the two proteins. Catalytic residues and the amides of the oxyanion hole are shown in sticks.

The second region for mutagenesis involves removing correlated motions from HNL3V. The residues 104-106 in HNL3V are located at the loop connecting the β-sheet β6 and the beginning of the lid domain. In HNL3V, Asn104 is directly connected to Trp203, whose movement is correlated to Thr204, Gln206, and the catalytic Asp207. The SPM of SABP2 shows no such correlations. Residues Ala105 and Ala106 (correspond to Asn104 and Ser105 in HNL3V) do not appear in the SPM indicating that their motions are not strongly correlated to any other residues. Residue Phe107 (corresponds to Val106 in HNL3V) appears in the SPM, but in contrast to HNL3V, its motion is not correlated with the catalytic aspartate. Therefore, the SPM predicted that substitutions Asn104Ala, Ser105Ala, Val106Phe in HNL3V would remove the connection of motions between this loop and the catalytic aspartate.

Four of the five SPM mutations predicted to fix the oxyanion hole orientation and the correlated motions of the catalytic aspartate are in the second or third shell outside the active site (Fig. 4). Residues 104-106 are on a loop adjacent to the catalytic serine and histidine. The G176S substitution is on a loop adjacent to one of the oxyanion hole residues. The fifth of the five substitutions is in the active site since it replaces the oxyanion hole residue Cys81 with leucine. Variant HNL8V contains the HNL3V substitutions and all five substitutions predicted by the SPM. Variant HNL7V contains the HNL3V substitutions and only four of the substitutions predicted by the SPM; the Ser105Ala substitution is omitted because the similarity of serine and alanine suggested that this substitution may be less important.

**Fig. 4.**
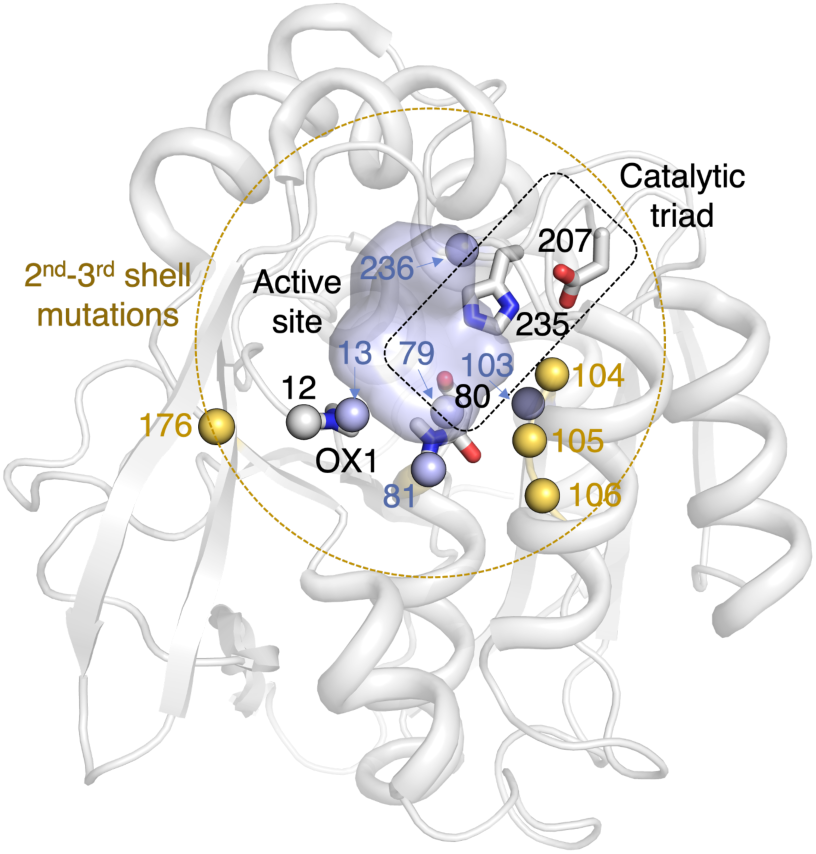
Location of the five substitutions (C81L N104A, S105A V106F G176S) added to HNL3V to create HNL8V. The catalytic triad residues (Ser80, Asp207, His235) and the oxyanion hole residues (Ile12, Cys81Leu) are blue spheres at the Cɑ with the side chains shown as sticks. The initial set of three substitutions (blue spheres at the Cɑ) were next to catalytic residues: Thr11Gly next to Ile12, Glu79His next to Ser80, Lys236Met next to His235. Four of the substitutions predicted by the SPM (yellow) are 2^nd^ and 3^rd^ shell changes outside the loops holding the catalytic residues: Gly176Ser outside Ile12, Asn104Ala, Ser105Ala, Val106Phe outside Ser80 and His235. One of the substitutions predicted by the SPM was the replacement of a catalytic residue (Cys81Leu, OX2) next to the catalytic serine (Ser80). Stabilizing substitution His103Val is shown as a blue sphere at the Cɑ. Part of the main chain trace is not shown for clarity; in reality the active site is buried.

### Designed substitutions yield an esterase more efficient than SABP2

The predicted variants HNL8V (HNL3V plus the five predicted SPM substitutions) and HNL7V (omit the Ser105Ala substitution from HNL8V) both proved to be good esterases, Fig. 5, Table 1. HNL7V was 50-fold more catalytically efficient (k_cat_/K_M_ of 25,000 M^-1^ min^-1^) than HNL3V and 290-fold more efficient than *Hb*HNL. HNL8V showed a slightly lower catalytic efficiency (k_cat_/K_M_ of 23,000 M^-1^ min^-1^), but the k_cat_ was 2-fold higher (k_cat_ of 8.1 ± 0.4 min^-1^). HNL6V, containing substitutions C81L, N104A, and G176S, showed a 2.4-fold improvement in turnover (k_cat_ of 0.59 ± 0.14 min^-1^) and a 100-fold improvement in binding (K_M_ of 0.03 ± 0.003 mM) relative to *Hb*HNL, resulting in a 240-fold improvement in catalytic efficiency (k_cat_/K_M_ of 20,000 *M*^-1^·min^-1^).

**Fig. 5.**
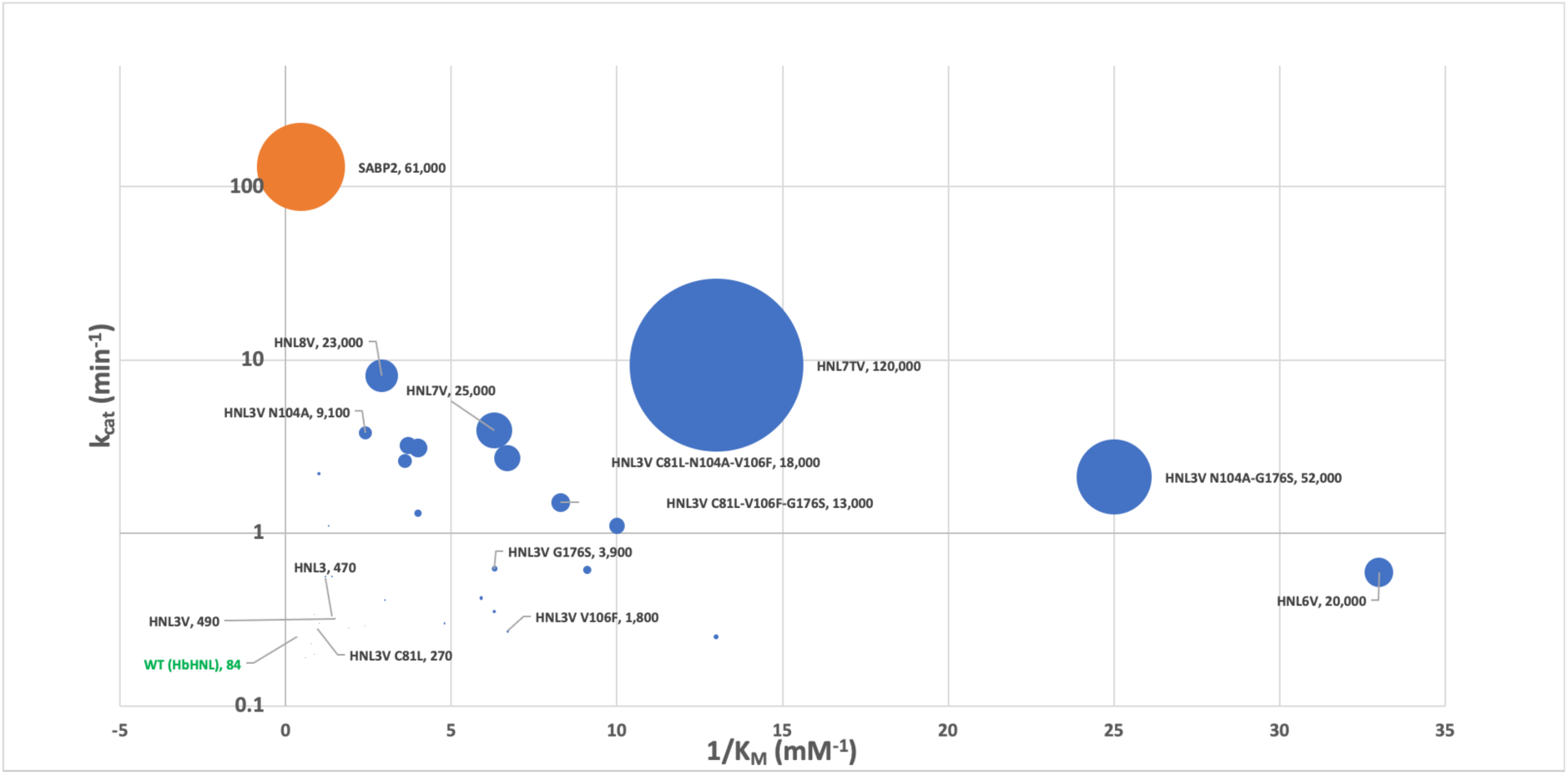
Protein engineering of *Hb*HNL (green text) for improved esterase activity (k_cat_/ K_M_ represented by ball size) yielded a variant (largest blue ball) with two-fold better catalytic efficiency than SABP2 (orange ball). Most variants showed better binding than SABP2, i.e. are further right on the x-axis, but the catalytic step was slower, i.e. lower on the y-axis. All *Hb*HNL variants are shown with blue balls. The k_cat_/K_M_ values in units of M^-1^ min^-1^ are shown for selected variants.

Of the four substitutions added to HNL3V to create HNL7V, the Asn104Ala substitution had the largest effect on catalytic activity. Excluding HNL7TV (discussed below), 9 of the 10 fastest variants and the 5 most efficient variants all contained N104A. HNL3V N104A’s turnover rate was as fast as HNL7V (k_cat_ of 3.8 ± 0.2 vs. 3.9 ± 0.4 min^-1^), though its catalytic efficiency (k_cat_/K_M_ of 9,100) was nearly 3-fold lower because of poorer binding (K_M_ of 0.42 ± 0.08 vs. 0.16 ± 0.07 for HNL7V). All of the intermediate variants between HNL3V N104A and HNL7V - those containing some combination of mutations C81L, N104A, V106F, and G176S - showed lower activity and poorer efficiency than HNL3V N104A and HNL7V. Mutations C81L, N104A, V106F, and G176S therefore interact epistatically and result in non-linear fitness effects. To confirm the importance of N104A for esterase catalysis, we made the reverse substitution in SABP2 to create SABP2 A105N and found a 4.7-fold decrease in enzyme turnover (k_cat_ dropped from 134 to 29 min^-1^). A decrease in binding (K_M_ increased from 2.2 to 3.0 m*M*) resulted in a 6.3-fold decrease in catalytic efficiency (k_cat_/K_M_ dropped from 61,000 to 9,600 *M*^-1^ min^-1^) as compared to SABP2. Relative to the best *Hb*HNL-based variant that does not contain N104A (HNL3V V106F-G176S), SABP2 A105N showed a 9-fold higher turnover rate. We expected a larger decrease in catalysis in SABP2 A105N given the importance of N104A for improved esterase activity in *Hb*HNL-based variants. This smaller than expected change suggests that epistasis plays a role in esterase activity for both SABP2- and *Hb*HNL-based variants.

The best variant, HNL7TV, contained eight substitutions and was 1390-fold more catalytically efficient (k_cat_/K_M_ of 120,000 M^-1^ min^-1^) than *Hb*HNL and two-fold more catalytically efficient than the benchmark esterase SABP2 (Fig. 5). We made the N104T substitution because although SABP2 contains Ala at position 105, Thr is the most highly conserved amino acid at that position among homologous esterases (Supplementary Fig. 3). Thus the N104T mutation was expected to yield soluble, active protein. HNL7TV showed the highest activity (k_cat_ of 9.3 ± 0.3) of all variants, a 37-fold improvement, and a 28-fold improvement in binding compared to *Hb*HNL (K_M_ of 0.08 ± 0.04 m*M*).

All of the *Hb*HNL variants containing SPM-predicted substitutions demonstrated an improvement in substrate binding and catalytic efficiency over *Hb*HNL. Notably, 65% and 59% of the variants showed at least 5-fold enhancements in binding over *Hb*HNL and SABP2, respectively. Despite the increase in catalytic activity, the fastest variants did not necessarily translate to superior binders; only three out of the ten fastest variants ranked in the top ten for binding and we found no correlation (R^2^ = 0.01) between k_cat_ and K_M_ (Supplementary Fig 4, Supplementary Table 2).

### Molecular dynamics simulations reveal restored oxyanion hole

Molecular dynamics (MD) simulations confirm a distorted OX1 N positioning in HNL3V as compared to SABP2 (Fig. 6a) similar to that seen in the x-ray structure comparisons of *Hb*HNL and SABP2 above (Fig. 2). Multiple replica nanosecond timescale MD simulations reveal a distribution in the distance between OX1 N and the serine Cɑ of ca. 5.5 Å in SABP2 as compared to 5.8 Å in the x-ray, but a much longer distance of ca. 6.2 Å for HNL3V as compared to 6.8 Å in the x-ray of *Hb*HNL. This shorter distance in SABP2 allows OX1 N to orient properly to stabilize the developing negative charge during catalysis.

**Fig. 6.**
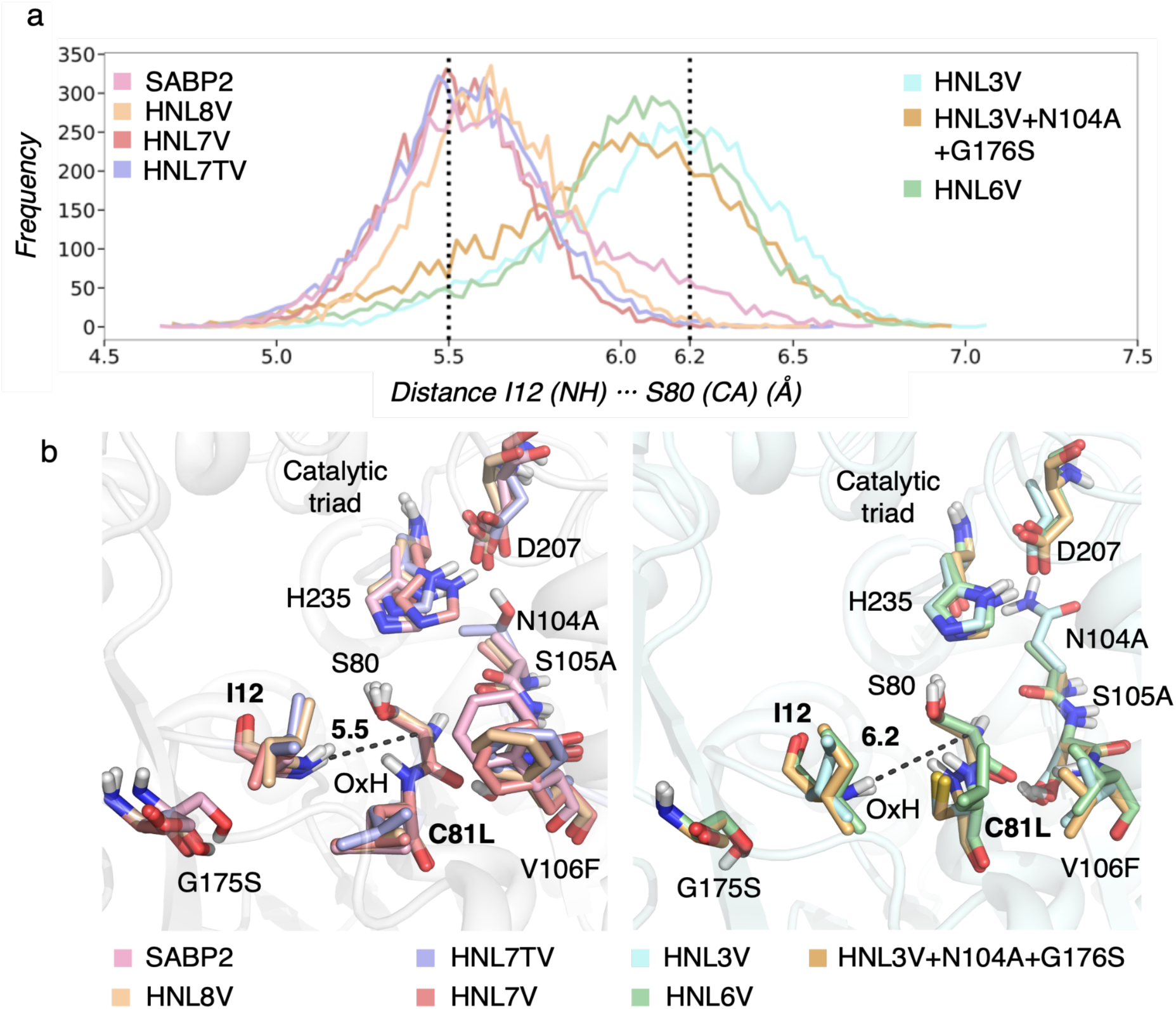
The OX1 N position of the most active variants mimic that of SABP2. **a**, Histogram of the distances (in Å) between the amide backbone of OX1 N (Ile12 or Ala13) and Cɑ of the catalytic serine (Ser80 or 81) for the following proteins: SABP2 (pink), HNL3V (blue), HNL3V N104A G176S (orange), HNL6V (green), HNL7V (dark pink), HNL7TV (purple), *Hb*HNL8V (brown). **b**, Overlay of most populated conformations of the variants presenting properly preorganized oxyanion hole residues (i.e., distances of ca. 5.5 Å, left panel): SABP2 (pink), HNL7V (dark pink), HNL7TV (purple), HNL8V (brown), and those presenting a non-optimal oxyanion hole positioning (i.e., distances of 6.2 Å): HNL3V (blue), HNL3V N104A G176S (orange), HNL6V (green). The labels refer to *Hb*HNL residue numbering.

MD simulations of many variants displaying higher levels of esterase activity matched the OX1 N positioning in SABP2, while those with lower esterase activity matched the OX1 N positioning in HNL3V (Fig. 6). An overlay of the most populated conformations in the multiple replica nanosecond timescale MD simulations confirm a restored oxyanion hole similar to that found in SABP2 for HNL7V, HNL7TV, and HNL8V. HNL3V and HNL6V do not present a catalytically productive positioning of the oxyanion hole residues, in line with their inferior esterase catalytic efficiency.

However, the OX1 N positioning in the MD simulation does not match the esterase activity in several cases. The second most efficient variant, i.e., HNL3V N104A-G176S, adopts longer distances between the catalytic serine and the oxyanion hole residue similar to those found in the less efficient HNL3V and HNL6V. Similarly, HNL7V and HNL7TV show similar OX1 N positioning, but their esterase activity differs. Thus, the properly restored oxyanion hole explains only part of the enhancements in esterase activity.

### MD simulations identify changes in the p*K*_a_ of catalytic aspartate

One of the substitutions that enhanced esterase catalytic efficiency is N104A/T, which lies close to the catalytic histidine and aspartate. The electrostatic environment of the catalytic triad has a profound effect on the catalytic activity of cysteine and serine proteases.^[24, 25]^ The catalytic Asp in the serine peptidase trypsin and a cysteine peptidase of the papain superfamily lie in different electrostatic environments creating different p*K*_a_ values for the Asp-His of the triad. MD simulations of variants with and without the N104A substitution show similar positions of OX1 N indicating that this substitution does not affect the oxyanion hole reorganization.

We hypothesize that the close location of N104A/T to the catalytic aspartate alters its local environment and p*K*_a_ value. We estimated the p*K*_a_ of Asp using the deep learning approach pKaI^[26]^ at each frame of an MD simulation and validated our predictions using constant pH MD simulations^[25]^ (Fig. 7). The catalytic aspartate in SABP2 was highly flexible causing changes to the local environment of aspartate and its p*K*_a_. The predicted p*K*_a_ values for the catalytic aspartate varied between 2 and 7. For the HNL variants containing the N104A/T substitution, the pKaI predictions match constant pH MD simulations, both predicting a p*K*_a_ of ∼5 for the catalytic aspartate. The catalytic aspartate is less flexible in these variants and the range of predicted p*K*_a_ values is narrower than that for SABP2. In contrast, HNL3V, which lacks the N104A substitution, has an estimated p*K*_a_ of <3. This shift in pKa is consistent with the altered solvation and the lower catalytic activity. The catalytic aspartate maintained its hydrogen bond to the catalytic histidine throughout all the simulations in all cases.

**Fig. 7.**
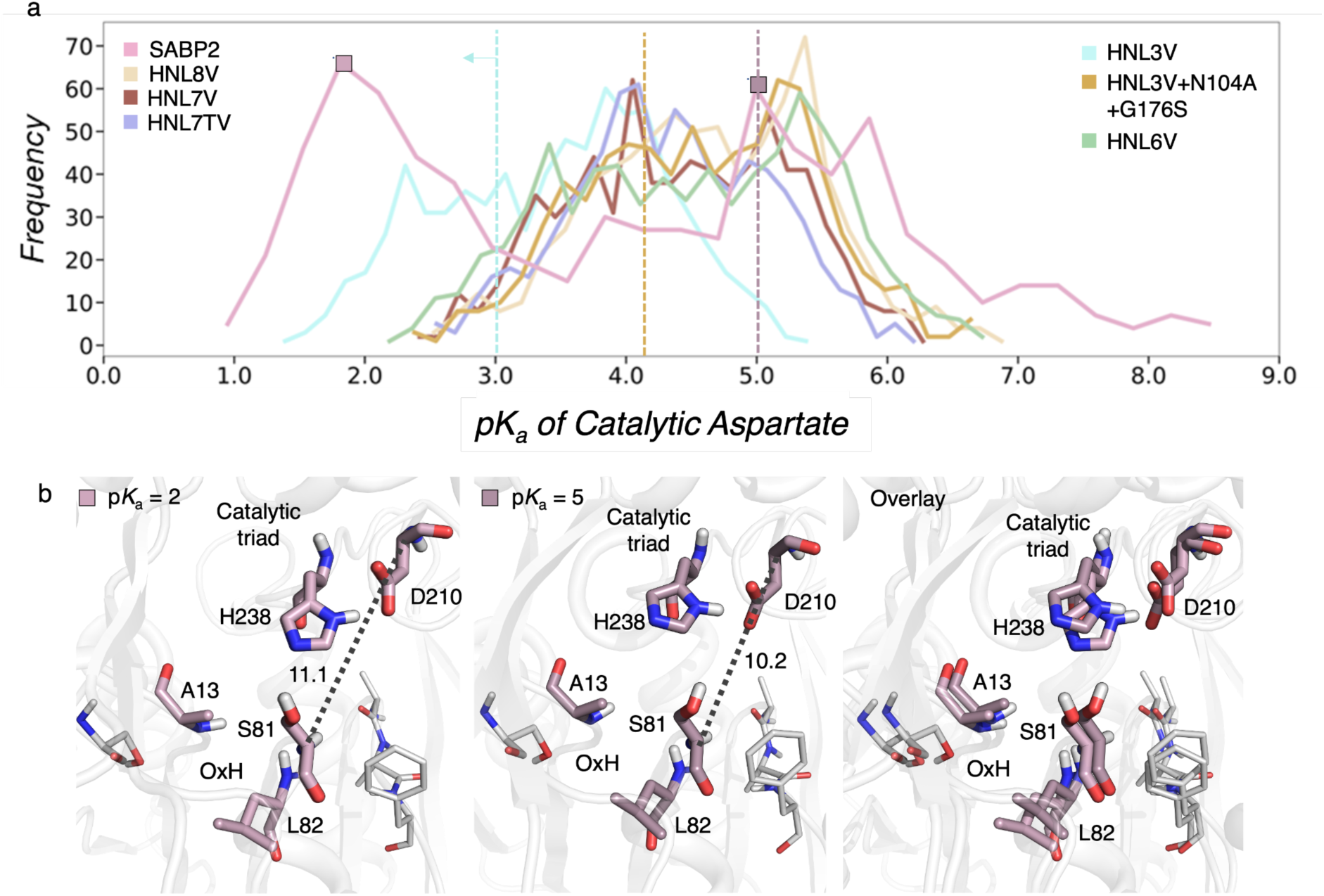
Estimation of the p*K*_a_ of the catalytic aspartate in SABP2 (pink) and HNL variants. **a**. Histogram of the p*K*_a_ values of the catalytic aspartate as predicted by pKaI at multiple frames of the MD simulation of SABP2 (pink), HNL3V (cyan), HNL3V N104A G176S (gold), HNL6V (green), HNL7V (brown), HNL7TV (purple), and HNL8V (light gold). The p*K*_a_ values estimated by constant pH MD simulations are marked with a vertical line for SABP2 (pink), HNL3V (cyan), and HNL3V N104A G176S (gold). b. Representative SABP2 conformation presenting a low p*K*_a_ value of ca. 2 (left panel), higher p*K*_a_ of ca. 5 (middle panel), and overlay of both conformations (right panel). The distances between the carbon alpha of the catalytic D210 and S81 are shown in Å.

### X-ray structure of HNL6V reveals aspartate hydrogen bond network

To confirm the changes in OX1 N positioning and solvation of the catalytic aspartate, we solved the x-ray crystal structure of HNL6V, which contains three of the five substitutions that were added to HNL3V to create HNL8V. Substitutions C81L, N104A, and G176S are present in HN6V, but substitutions V106F and S105A are missing.

The structure of HNL6V aligns closely with the structure of wild-type *Hb*HNL. The catalytic domain (residues 1-114, 179-264) adopts the ɑ/β-hydrolase fold and contains the catalytic triad and oxyanion hole residues. The lid or cap domain (residues 115-178) covers the active site to create a substrate binding pocket. The seven amino acid substitutions in HNL6V were in and around the active site, and none of the residues were on the outer protein surface. Six of the substitutions are in the catalytic domain; only Gly176Ser is in the lid domain.

The OX1 N in HNL6V has moved closer to the position of SABP2, but remains intermediate between *Hb*HNL and SABP2 (Fig. 8). The distance from OX1 N in HNL6V and OX1 N in SABP2 is 0.9 Å, while the corresponding distance between the three *Hb*HNL structures and SABP2 is longer: 1.2 ± 0.2 Å. The internal distance between OX1 N and serine Cɑ is 6.2 Å in HNL6V, which is intermediate between that in SABP2 (5.8 Å) and the three *Hb*HNL structures (6.8 ± 0.2 Å), see Fig. 2 above. Thus, the substitutions in HNL6V moved the main chain nitrogen (OX1 N) closer to its position in SABP2.

**Fig. 8.**
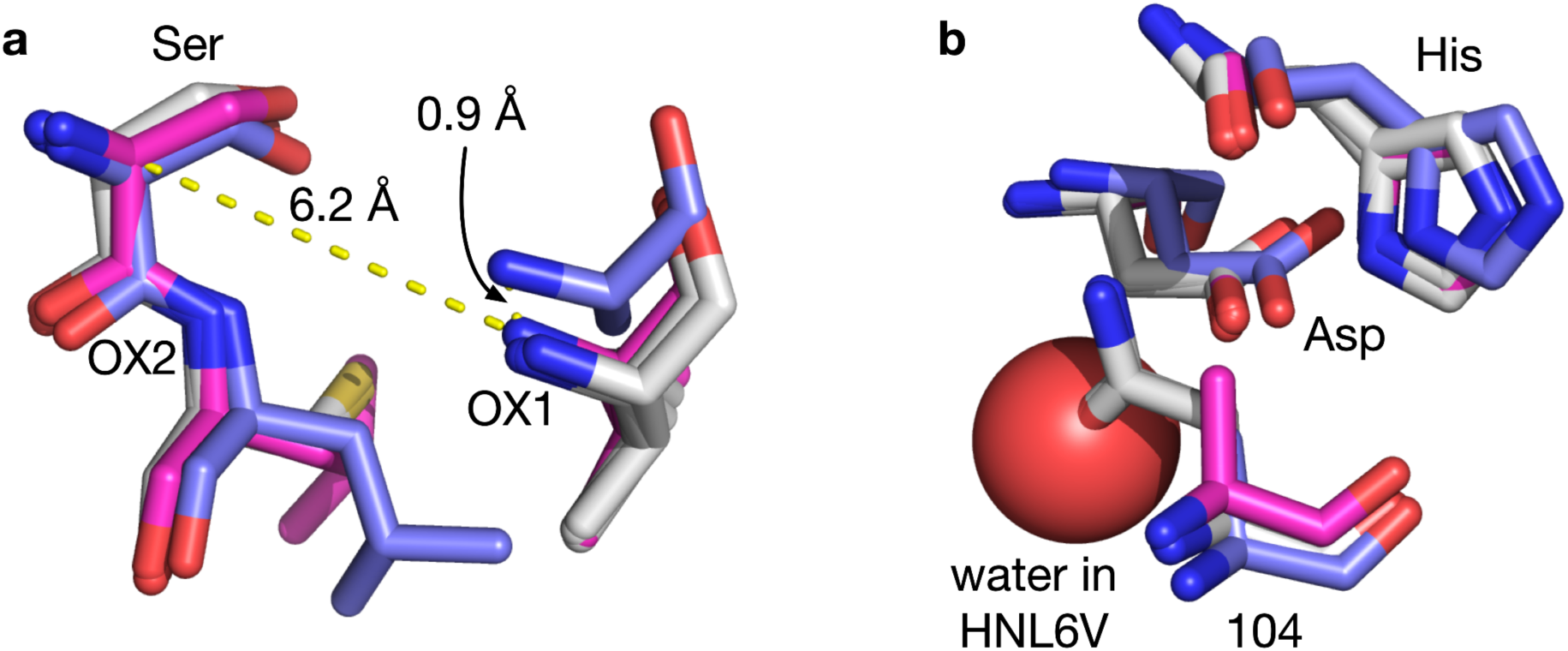
Changes in (a) the position of OX1 N and (b) the solvation of the catalytic aspartate in the x-ray structure of HNL6V (magenta carbons) as compared SABP2 (blue carbons) or *Hb*HNL (three structures, white carbons). **a)** The position of OX1 N in HNL6V has moved closer to the corresponding position in SABP2. The distance between OX1 N in SABP2 and HNL6V (0.9 Å) is shorter than the distance between SABP2 and the three *Hb*HNL structures (1.2 ± 0.2 Å). The internal distance between OX1 N and serine Cɑ is 6.2 Å in HNL6V, which is intermediate between that in SABP2 (5.8 Å) and the three *Hb*HNL structures (6.8 ± 0.2 Å). **b)** The Asn104Ala substitution in HNL6V creates space for a water molecule (red sphere) near the catalytic aspartate. This space is blocked in *Hb*HNL by the side chain of the asparagine. The alignment of the structures minimized the RMSD of the corresponding Cɑ atoms in the entire protein.

The Asn104Ala substitution in HNL6V created space for a water molecule that hydrogen bonds to the catalytic aspartate. In the *Hb*HNL structures, the asparagine residue fills this region, but does not interact with the catalytic aspartate. Upon replacement of Asn104 with alanine, more space is available. A water occupies this region in the x-ray structure of HNL6V and contributes a hydrogen bond to the catalytic aspartate (O-O distance is 3.1 Å).

## Discussion

Previous computational designs have required additional experimental optimization to reach catalytic efficiencies comparable to Nature’s enzymes. Computational design of a luciferase using Rosetta combined with deep learning yielded impressive catalytic efficiencies of 10^6^ M^-1^·s^-1^ but this design also included experimental optimization of the ligand-binding pocket.^[27]^ A bioinformatics-based design of a hydroxynitrile lyase from an esterase yielded a catalytic efficiency of 3,300 M^-1^·s^-1^.^[28]^ It required >120 substitutions, which make it difficult to explain how each substitution contributes to catalysis. The computational design of multi-step reactions has been less successful. A designed retroaldolase showed a k_cat_/K_M_ of 0.2 M^-1^·s^-1^ and reached a k_cat_/K_M_ of 34,000 M^-1^·s^-1^ only after multiple rounds of directed evolution.^[29, 30]^ Designed esterases based on a designed cysteine-histidine dyad and oxyanion hole yielded catalytic efficiencies of k_cat_/K_M_ of 10-400 M^-1^·s^-1^.^[12]^ One attempt at esterase design with serine-histidine-aspartate catalytic triads failed to complete a catalytic cycle. The enzymes could only react irreversibly with fluorophosphonate probes.^[9]^ A more recently designed esterase showed a catalytic activity 1000-times lower than commercially available esterases, but saturation kinetics were not reported.^[8]^ A Kemp eliminase also required eight substitutions outside the active site for high catalytic efficiency, but these were not rationally predicted, but found with experimental directed evolution.^[31]^ This inability to design efficient enzymes limits new applications of enzymes in medicines, non-polluting manufacture of fine chemicals and pharmaceuticals, food processing, and biodegradation of environmental contaminants.

The low catalytic efficiencies achieved by computational enzyme design have been associated with non-optimal arrangements of the catalytic residues for transition state(s) stabilization, the lack of a proper description of the conformational changes key for substrate binding and product release, and the limitation of introducing mutations in the active site pocket only.^[1]^ Our approach of identifying the correlated motions established by catalytic residues with second and third shell mutations has achieved catalytic efficiencies surpassing that of the reference SABP2 enzyme.

Comparison of the shortest path maps for HNL3V and SABP2 revealed differences in correlated movements in the two enzymes. To engineer increased esterase activity into HNL3V, we focused on correlated movements connected to the active site residues. HNL3V contained one movement associated with catalytic aspartate that was missing from SABP2. We hypothesized that this movement should be removed from HNL3V to increase esterase activity. HNL3V also lacked a correlated movement associated with an oxyanion hole residue that was present in SABP2. We hypothesized that this movement should be added to HNL3V to increase esterase activity. To add or remove movements, we replaced residues in HNL3V with the corresponding residues from SABP2. Only residues within the SPM of either HNL3V or SABP2 were changed. Since HNL3V and SABP2 share 45% sequence identity, only five amino acid substitutions were required. Four of the five substitutions were outside of the active site demonstrating that the shortest path maps identify residues outside the active site that contribute to catalysis.

The resulting variant, HNL8V, showed a 25-fold increase in both catalytic rate (k_cat_) and a 1.9-fold improvement in K_M_ for esterase catalysis demonstrating the value of the SPM-based predictions. The interactions between the substitutions showed negative cooperativity with respect to K_M_, but positive cooperativity with respect to k_cat_. Individually, the five substitutions showed modest improvements in K_M_ (mean of 2.5 ± 1.6-fold improvement). If the improvements act additively, then HNL8V should show a 34-fold improvement in K_M_, but it showed only a 1.9-fold improvement. The eighteen-fold lower observed value indicates negative cooperativity between the five substitutions with respect to K_M_. For k_cat_, four of the substitutions showed modest changes, but N104A showed a twelve-fold improvement (mean = 3.2 ± 4.9-fold improvement). If the improvements act additively, then HNL8V should show a 8.1-fold improvement in k_cat_, but it showed a 25-fold improvement. The 3.1-fold higher observed value indicates positive cooperativity between the five substitutions with respect to k_cat_.

This cooperativity is consistent with the notion that cooperative movements cause the changes in esterase activity. The predicted SPM substitutions – C81L, V106F, and G176S – interact epistatically, Fig. 9. The effect of all three substitutions combined is more than twice the effect of the sum of the three individual substitutions. The conformational changes induced by each substitution suggest a mechanism for this epistasis. The replacement of a glycine by a serine at position 176 changes the backbone conformation, which properly positions the oxyanion hole residue Ile12 as (OX1)-Ala13 in SABP2. Fixing the orientation of the other oxyanion hole residue Cys81 (OX2-Leu82 in SABP2) requires both Cys81Leu and Val106Phe. The side chain of Leu81 can adopt two different conformations in HNL3V; one conformation hinders catalysis as it blocks access of the ester substrate to the active site pocket. Mutation Val106Phe restricts the side chain of Leu81 to the conformation that allows ester binding for catalysis. As noted in the results section, the effect of the mutation N104A/T is not connected to the oxyanion hole reorganization, but rather to the change of the electrostatic environment of the catalytic aspartate. Variants containing the key mutations for fixing the oxyanion hole (C81L, V106F, and G176S) together with N104A/T show the highest esterase catalytic efficiency.

**Figure 9.**
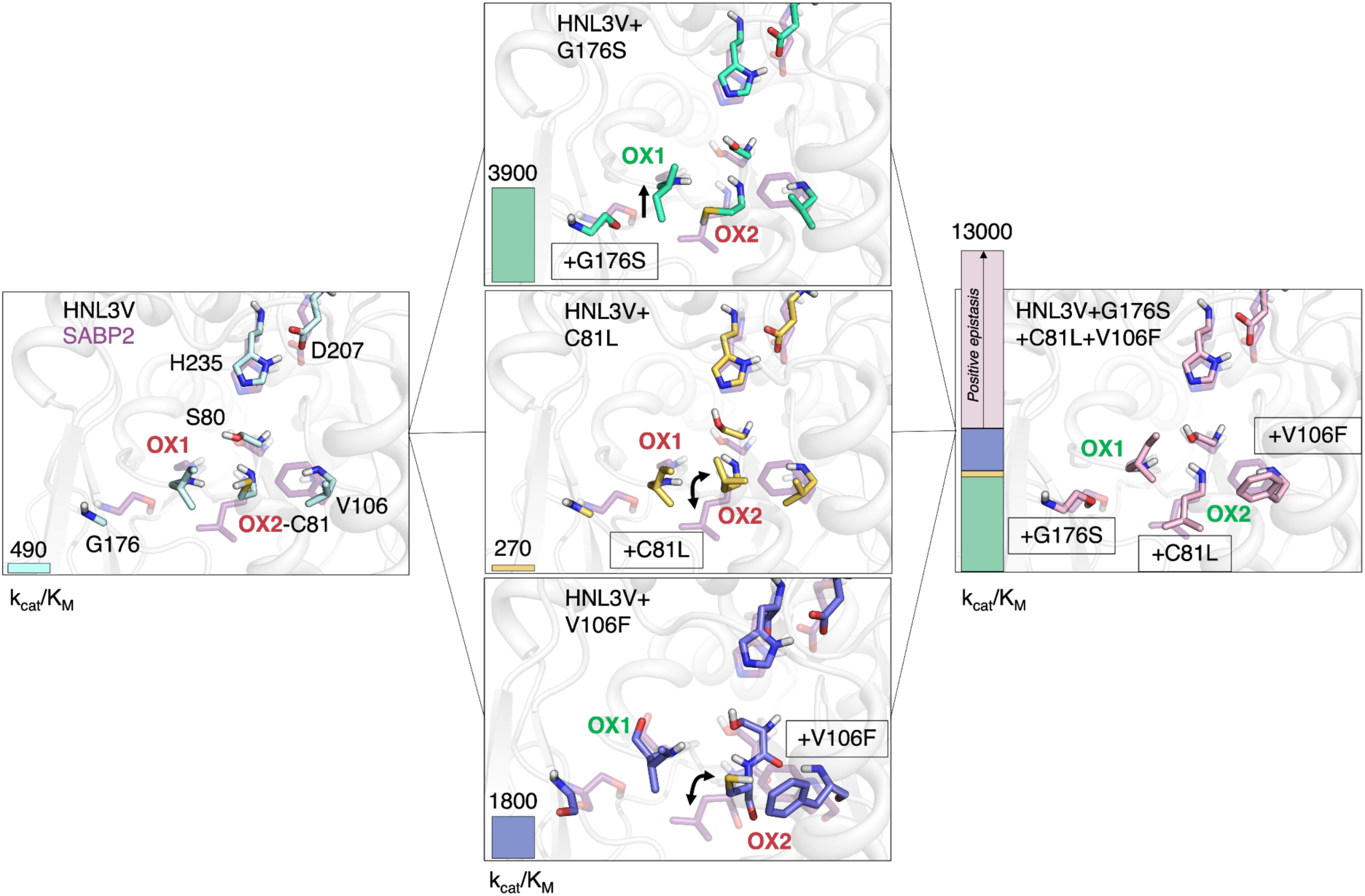
Substitutions that enhance catalytic activity act cooperatively. Representative conformations for the HNL variants: starting variant HNL3V (left panel, light blue carbons), the singly mutated variants HNL3V+G176S (center panel, green carbons), HNL3V+C81L (gold carbons), HNL3V+V106F (dark blue carbons), and the triple variant (right panel, pink carbons). All panels include a representative conformation of the active site and oxyanion hole residues of SABP2 (purple carbons) and a bar at the left side indicating the catalytic efficiency (k_cat_/K_M_, M^-1^·min^-1^) of each HNL variant. The combined effect of the three individual mutations is higher than their sum, due to positive sign epistasis highlighted in pink and with an arrow. The color of the labels of the oxyanion hole residues (OX1, OX2) indicate a proper (green) or bad (red) orientation as compared to SABP2. Double arrows mark the different conformations of the sidechain of C81 with respect to SABP2.

Combining the SPM analysis with a multiple sequence alignment of esterases yielded the best variant, HNL7TV. The substitution N104A yielded the largest increase in k_cat_ of all the substitutions tested. The multiple sequence alignment showed that most esterases contained threonine, not an alanine, at this position. The substitution N104T yielded the best variant with approximately twice the catalytic efficiency (k_cat_/K_M_) of SABP2.

Although the shortest path maps combined with sequence comparison identified the substitutions needed to improve catalytic activity, they did not identify why the substitutions increased catalytic activity. Gaining insight on why esterase activity increased required additional experiments combined with computational modeling. Molecular dynamics simulations support the notion that substitutions have repositioned the main chain of the oxyanion hole residue, Ile12. The 25-fold increase in k_cat_ from HNL3V vs. HNL7TV is consistent with previous experiments that disrupted the oxyanion hole in enzymes using site-directed mutagenesis. Removing an N-H from the oxyanion hole in subtilisin lowered k_cat_ approximately 100-fold.^[32, 33]^ Experiments with ketosteroid isomerase^[34]^ and a decarboxylase^[35]^ gave similar estimates for the contributions of an oxyanion hole N-H to catalysis.

Both molecular dynamics simulations and an x-ray structure of HNL6V show changes in the local environment of the catalytic aspartate that might impact its p*K*_a_. The mutation of the catalytic aspartate in trypsin by an asparagine lowers the p*K*_a_ of histidine by 1.5 pH units and dramatically reduces the catalytic activity.^[36, 37]^ One mechanism to explain the role of the catalytic aspartate in serine proteases is the formation of a low-barrier hydrogen bond between His-Asp, especially in the transition state or in enzyme-intermediate complexes.^[38, 39]^ This mechanism requires that both histidine and aspartate present similar p*K*_a_ values, which for aspartate was estimated to be around 6.7.^[40]^ The increase in catalytic efficiency observed for the variants containing N104A/T directly in contact with the catalytic aspartate suggest that this mutation impacts the p*K*_a_ of the catalytic aspartate. Our calculations predict higher p*K*_a_ values for aspartate especially in SABP2, and in the most efficient variants HNL7V, HNL7TV and HNL8V. This finding is in line with the requirement of matching p*K*_a_ values between His-Asp for the formation of a low-barrier hydrogen bond at the transition states and/or tetrahedral intermediates formed along the multistep esterase mechanism.

There must be additional contributions to esterase catalysis in SABP2 besides those identified here. The k_cat_ of the best variant, HNL7TV, is still about thirteen-fold lower than that for SABP2 indicating that additional substitutions are needed to fully match the k_cat_ of SABP2. Even more distant substitutions are likely required to modulate the conformational landscape of the enzymes and generate a SABP2-like environment of the catalytic His-Asp dyad for esterase catalysis. The higher flexibility of the catalytic aspartate in SABP2 identified in our MD simulations may contribute to its higher k_cat_.

## Methods

Chemicals were purchased from commercial suppliers and used without further purification.

### Site-directed mutagenesis to create enzyme variants

The gene encoding the wild-type HNL from *Hevea brasiliensis* in the pSE420 plasmid^[41]^ was recloned into a pET21a(+) plasmid. *Hb*HNL enzyme variants were constructed via inverse PCR^[42]^ using non-overlapping mutagenic primers in a sequential manner (Supplementary Table 3). Briefly, mutagenic primers anneal to the plasmid template in a back-to-back, outward-facing orientation and are amplified using New England Biolabs (NEB) Q5 HiFi polymerase (M0491S) to produce a linear, double-stranded DNA product containing the desired mutation(s). Primers were designed using the NEBaseChanger (https://nebasechanger.neb.com) web tool and checked for secondary structure and self-dimer/ heterodimer propensity with IDT’s OligoAnalyzer Tool (https://www.idtdna.com/pages/tools/oligoanalyzer). Primers were purchased from Integrated DNA Technologies (Coralville, IA) and used without further purification. PCR was performed using a BioRad 2000 Thermal Cycler with the following conditions: initial denaturation at 98 ℃ for 30 sec, 30 cycles of denaturation (98 ℃ for 30 sec), annealing (calculated annealing temperature for 25 sec), and extension (72 ℃ for 150 sec), and a final extension step of 72 ℃ for 2 minutes. The PCR products were treated with a KLD enzyme mix (NEB M0554S), which phosphorylates the 5’ ends of the linearized PCR products, ligates the phosphorylated ends, and degrades the original plasmid template. Five µl of the KLD product was used directly to transform chemically-competent *Escherichia coli* DH5ɑ cells (NEB C2988) according to the manufacturer’s protocol and plated on lysogeny broth (LB) plates containing 100 µg/ml carbenicillin. After overnight growth at 37 °C, individual colonies were picked and grown up overnight in LB media, and the plasmids were extracted via NEB Monarch Plasmid mini-prep kit (NEB T1010). Plasmid concentrations were measured spectrophotometrically at 260 nm via a Nanodrop 2000 (Thermo Scientific) and diluted to <1.0 OD units if necessary. Sanger sequencing from Genewiz/Azenta Life Sciences was used to confirm mutations. The sequence-confirmed plasmid was transformed into *Escherichia coli* strain BL21(DE3) chemically competent cells (NEB C2527) according to NEB’s transformation protocol and plated on LB plates containing 100 µg/ml carbenicillin.

SABP2 A104N was constructed via isothermal assembly (also called Gibson assembly)^[43]^ using a gene fragment ordered from Twist Biosciences (San Francisco, CA). We used the NEB Gibson Assembly® Master Mix kit to perform the assembly under the following conditions: template DNA (pET21a(+) plasmid containing SABP2 gene), enzyme master mix, and the synthesized gene fragment were incubated in a thermocycler for 15 minutes at 50°C. 2 µl of the assembly reaction mixture was used directly for transformation into NEB 5-alpha competent *E. coli* included in the assembly kit. All subsequent steps, i.e. expression, purification, and assays, are as described above. For detailed information on the protocol, including primer design, please see NEB’s Gibson Assembly® Application Overview website (https://www.neb.com/applications/cloning-and-synthetic-biology/dna-assembly-and-cloning/gibson-assembly).

### Protein expression and purification

LB media containing carbenicillin (100 µg/ml, 5 ml) was inoculated with a single bacterial colony from an agar plate and incubated in an orbital shaker at 37 °C and 240 rpm for 15 h to create a seed culture. A 1-L baffled flask containing terrific broth-amp media (250 ml) was inoculated with 2.5 mL of seed culture. The pre-induction culture was incubated at 37 °C and 240 rpm for 3–4 h until the absorbance at 600 nm reached 0.4-1.0. The culture was then transferred to ice for 30 minutes to cool. Isopropyl β-D-1-thiogalactopyranoside (0.75–1.0 m*M* final concentration) was added to induce protein expression, and cultivation was continued for 20-24 h at 18℃. The cells were harvested by centrifugation (7000 rpm, 15 min at 4 °C), resuspended in NiNTA loading buffer (10 m*M* imidazole, 50 m*M* Tris pH 8.0, 500 m*M* NaCl, 4 ml/g of wet cells), and either directly sonicated or frozen for storage and later purification. Cells were flash frozen in liquid nitrogen or a dry ice-ethanol bath and stored at -80°C. Frozen cells were thawed at room temperature or in a room temperature water bath, and fresh/thawed cells were disrupted by sonication (400 W, 40% amplitude for 3 min). The cell lysate was centrifuged to pellet the cell debris (4 °C, 20,000 rcf for 20 min) and the supernatant was mixed with 1-2.5 ml of NiNTA resin (pre-equilibrated with 10 ml of NiNTA loading buffer) and incubated for 45 minutes at 4 °C with rotation (10 rpm). The resin/ supernatant mixture was loaded onto a 25 ml column (Bio-Rad) and the resin was washed with 10 column volumes each of buffer containing increasing amounts of imidazole (25-50 m*M* imidazole, 50 m*M* Tris pH 8.0, 500 m*M* NaCl). The His-tagged protein was eluted with 10 column volumes of elution buffer (125 m*M* imidazole, 50 m*M* Tris pH 8.0, 500 m*M* NaCl) and collected in 1 ml fractions. The protein concentration of each elution fraction was determined from spectrophotometric measurements at 280 nm via Nanodrop 2000 (Thermo Scientific). The calculated extinction coefficient was determined using the ProtParam web tool (https://web.expasy.org/protparam/). Protein gels were used to check for the presence and purity of protein and run using sodium dodecyl sulfate polyacrylamide gradient gels (NuPage 4−12% Bis-Tris gel from Invitrogen) using the Precision Plus Dual Color protein standard (BioRad, 5 µl/lane), run for 50 min at 120V, stained with SimplyBlue Safe Stain (Thermo Fisher Scientific), and destained 2x with milliQ UltraPure H2O. SDS-PAGE indicated a molecular weight of ∼30 kDa in agreement with the predicted weight of 31.1 kDa. The imidazole-containing elution buffer was exchanged by addition of BES buffer (5 m*M N*, *N*-bis(2-hydroxyethyl)-2-aminoethanesulfonic acid, pH 7.2, 14 ml) followed by ultrafiltration (Amicon 15-ml ultrafiltration centrifuge filter, 10 kDa cutoff) to reduce the volume to ∼ 250 µl. This addition of buffer and filtration was repeated four times. A 250-ml culture typically yielded 2-5 mg of protein.

### Enzyme assays

Enzyme activity was monitored at room temperature (typically 22±2 °C) in triplicate for 10 min using a SpectraMax 384 Plus microplate reader. Ester hydrolysis activity was measured at 405 nm using *p*-nitrophenyl acetate (pNPAc), which releases the yellow *p*-nitrophenoxide. The reaction mixture (100 µL; path length 0.29 cm) contained 0.01-7.0 mM pNPAc, 6–8% v/ v acetonitrile, 5 mM BES buffer, pH 7.2, and up to 15 µg enzyme. The slope of increase in absorbance versus time was measured in triplicate, fit to a line using linear regression, and corrected for spontaneous hydrolysis of pNPAc with blank reactions lacking protein, also measured in triplicate. The extinction coefficient used for calculations (ε_405_ nm = 11,588 cm^-1^ M^-1^) accounts for the incomplete ionization of *p*-nitrophenol at pH 7.2. For steady-state kinetic measurements, the enzyme concentration was determined by average absorbance at 280 nm measured in duplicate and normalized by subtracting a buffer blank. The enzyme concentrations in the assay solution ranged from 50 nM to 5 µM. k_cat_ and K_M_ were determined using a non-linear fit of the experimental data to the Michaelis-Menten equation using the solver program in Microsoft Excel or using the statistical program R (Huitema & Horsman, 2019).^[44]^

An alternative assay protocol (the KP protocol) results in faster rates relative to the BES protocol described in the previous paragraph. The reaction mixture volume, substrate concentrations, amount of enzyme, and absorption wavelength are the same between both assays. In the KP protocol, the substrate is dissolved in methanol instead of acetonitrile and uses 100 mM KP buffer, pH 7.5, 1% v/v acetonitrile, and an extinction coefficient (ε_405_ nm = 12,300 cm^-1^ M^-1^) that accounts for the effect of the change in pH on absorbance.^[45]^ The faster rate is due to the differing organic solvents; increasing the acetonitrile concentration decreases the observed rate, as has been previously described.^[46]^ All kinetic parameters reported in this manuscript were obtained using the BES protocol.

### Molecular modeling system preparation

The starting structures for the different enzymes (*HbHNL, HNL3V, HNL6V, HNL7V, HNL7TV, HNL8V, SABP2*) were generated with the predictions of the neural network AlphaFold 2 approach.^[47]^ The structures were prepared using the Python packages MDTraj,^[48]^ pytraj^[49]^ which is part of the cpptraj package,^[50]^ MDAnalysis,^[51]^ PyEMMA,^[52]^ and networkx.^[53]^

### Molecular dynamics simulation

The protocol applied for the MD equilibration phase was the one described by Roe and Brooks with small differences fine-tuned to our systems.^[54]^ For non-minimization steps, the bonds involving hydrogen are constrained by the SHAKE algorithm. Long-range electrostatic effects were modeled using the particle mesh-Ewald method.^[55]^ A 10 Å cut-off was applied to Lennard–Jones and electrostatic interactions. The MD protocol starts with the minimization phase of 1500 steps steepest descent method followed by 3500 steps of the conjugate gradient method with a positional restrain (*i.e.*, force constant of 5.0 kcal·mol^-1^·Å^-2^) to the protein heavy atoms. Then, a heating phase is performed with increasing the temperature from 25 K to 300 K during 20 ps of MD simulation, a Langevin thermostat with a collision frequency of 5 ps^-1^, and a positional restrain (i.e., force constant of 5.0 kcal·mol^-1^·Å^-2^) to the protein heavy atoms. The next step is the minimization and heating of all the atoms in the system. Starting with two minimization stages of 1000 steps steepest descent method followed by 1500 steps of the conjugate gradient method each with a positional restrain (i.e., force constant of 2.0 kcal·mol^-1^·Å^-2^ in the first minimization and 0.1 kcal·mol^-1^·Å^-2^ in the second) to the protein heavy atoms. Then, a third minimization phase of 1500 steps steepest descent method followed by 3500 steps of the conjugate gradient method without any positional restraint is performed. Afterwards, the system is heated in the same way as previously defined. Finally, a five-round equilibration phase at the NPT ensemble with a constant pressure of 1 atm is performed: whereas the first four were done with the Berendsen barostat, the fifth one with Monte-Carlo barostat. Langevin thermostat with a collision frequency of 1 ps^-1^ was used in the five equilibration rounds. The first two equilibration rounds of 5 ps had a positional restraint to the protein-heavy atoms with a force constant of 1.0 and 0.5 kcal·mol^-1^·Å^-2^, respectively. A third round of 10 ps equilibration is followed with positional restraint to the backbone-heavy atoms with a force constant of 0.5 kcal·mol^-1^·Å^-2^. The fourth equilibration round of 10 ps was performed without any restraint. The last equilibration round was of 1 ns without any restraint. The production runs were performed at the NVT ensemble with the Langevin thermostat with a collision frequency of 1 ps^-1^ during 250 ns. Finally, three replicas of equilibration and production runs were performed for each homodimer, reaching a total simulation time of 750 ns for *HbHNL, HNL3V, HNL3V_104A_176S, HNL6V, HNL7V, HNL8V, SABP2* systems, respectively. The MD trajectories were analyzed using the Python packages MDTraj,^[48]^ pytraj^[49]^ which is part of the cpptraj package,^[50]^ MDAnalysis,^[51]^ PyEMMA,^[52]^ and networkx.^[53]^

### Constant pH Molecular dynamics simulation

Constant pH molecular dynamics simulations were done following the same protocol described in the molecular dynamics simulation section. Residues are allowed to change protonation state in the fifth equilibration and the production runs. All systems were simulated at pH values from 4.5 to 8.0 with a 0.5 spacing. The protonation state changes were attempted every 100 steps. The following 100 steps were used to relax the solvent after a successful attempt. A salt concentration of 0.1 was used. For SABP2 HID6, HID11, HID15, HIE32, HIE80, HIP113, HID158, ASP210, HID238, HID257 residues were selected to titrate, and for HNL variants HID5, HID10, HID14, HID20, HIE31, HIE79, HIP112, ASP207, HID235 were selected to titrate. Three replicas of equilibration and production run of 30 ns were performed for SABP2, HNL3V, HNL3V G176S N104A. p*K*_a_ values are estimated from the extrapolation of the sigmoidal function.

### Molecular dynamics analysis

*Shortest Path Map (SPM) calculations.* The Shortest Path Map (SPM) analysis was performed using the MD simulations of SABP2 and HNL3V. For SPM calculation, the inter-residue mean distance and correlation matrices computed along the MD simulations need to be computed. From both matrices a simplified graph is drawn, in which only those pairs of residues displaying a mean distance shorter than 6 Å along the MD simulation time are connected through a line. The edge connecting both residues is weighted to the Pearson correlation value (dij=-log |Cij|). Short lines will be drawn for those pairs of residues whose motions are more correlated. The generated graph is further simplified to identify the shortest path lengths. Following this strategy, those lines in the graph that are shorter, i.e. the connecting residues are more correlated, and that play a substantial role in the enzyme conformational dynamics are detected. The generated SPM graph is then drawn on the 3D structure of the enzyme. More details about our SPM tool can be found in references 1 and 5.

### X-ray crystal structure determination

HNL6V containing a C-terminal 6His tag was expressed from plasmid pET21a(+) in *Escherichia coli* BL21 (DE3). The protein was purified using nickel-affinity chromatography and concentrated to 9.3 mg/ml. Crystallographic screening was done using Phoenix crystallography dispenser from Art Robbins Instruments Inc. Sitting-drop vapor diffusion trays, the low profile INTELLI-PLATE® from Art Robbins Instruments Inc, were used for crystallization setup. All the setup and washing procedure was done through the Art Robbins Instruments software Phoenix. Each crystallization drop contained 0.1 µl protein sample (9.3 mg/ml protein) and 0.1 µl well solution. A total of 960 conditions were tested. Crystals appeared within one day from the Index HT screen from Hampton Research Inc, under the condition of 0.1 *M* Bis-Tris, pH 5.5, 2 *M* (NH_4_)_2_SO_4_, and grew to the full size of 0.35 mm in three days. Two distinct crystals formed (Supplementary Fig. 5). The robotic screening crystals proved sufficient to refine the model so additional crystal screening trays were not needed.

The structural dataset was collected on beamline 24 ID-C (NE-CAT) at the Advanced Photon Source, Argonne National Laboratory (Supplementary Table 4). Crystals were transferred into cryoprotectant solutions consisting of well solution components and increasing concentration of sodium malonate. The final concentration of sodium malonate in the cryoprotectant solution was 1.2 *M*. Harvested crystals were flash frozen in liquid nitrogen. The datasets were collected at an oscillation angle of 0.2°. The crystal belonged to space group *C*222_1_, with unit cell parameters a = 47.054, b =106.378, c = 128.396 Å and, one molecule per asymmetric unit (Supplementary Table 5). The reconstructed ancestral hydroxynitrile lyase (PDB ID: 5tdx^[20]^) with 75.85% sequence identity was used for molecular replacement, and refined to a 2.3 Å resolution model. This 2.3 Å initial model was used to aid in the refinement of the second and final model resulting in a 1.99 Å structure. HKL2000^[56]^ was used to process collected data and Phaser^[57]^ in Phenix^[58]^ was used for molecular replacement and refinement (Supplementary Table 6). Refinement modeling was performed using Coot.^[59]^ The structure was refined to R_work_ and R_free_ values of 0.1844 and 0.2376, respectively.

Structural refinement revealed 2F_O_-F_C_ (blue) and F_O_-F_C_ (red/green) electron density near the active site, located around the catalytic serine Oɣ (Supplementary Fig. 6). This density suggested the presence of a bound molecule, perhaps in multiple orientations. Placing water, glycerol, malonate, or sulfate in this region did not improve the R-work and R-free statistics, nor did these placements satisfy the density. Placing water around the catalytic Serine Oɣ in multiple orientations did not reduce the F_O_-F_C_ density, suggesting the structure still required the addition of one or more molecules, and perhaps in sub-100% occupancies. Adding glycerol in multiple locations proved unsatisfactory due to the increase in red F_O_-F_C_ around pieces of the molecule unable to fit in the 2F_O_-F_C_ density properly. Placing glycerol in varying locations with occupancies summing to 100% in attempts to solve this issue remained insufficient to satisfy the 2F_O_-F_C_ density. When adding malonate, negative interactions with nearby amino acid residues became unavoidable, no matter the position or orientation of the malonate. Additionally, experimental placement of malonate with varying occupancy proved unfulfilling to the 2F_O_-F_C_ density. Finally, the inability to find a placement of sulfate molecules that would fulfill the active site density without interaction with each other concluded attempts to place a ligand in the active site. As this electron density near catalytic serine Oɣ remains unmodeled, this model should be considered a putative structure. Further refinement of other *Hb*HNL structures could potentially aid in identifying the structure in the currently unresolved F_O_-F_C_ density active site. Zuegg and coworkers^[13]^ also observed unidentified electron density near the active site during refinement of an x-ray crystal structure of wild-type *Hb*HNL. The final model was deposited in the Research Collaboratory for Structural Bioinformatics Protein Data Bank (PDB ID: 8euo).

The three structures of wild-type hydroxynitrile lyase from *Hevea brasiliensis* without a bound ligand used for comparison have the following protein data bank IDs: 6yas,^[20]^ 3c6x, ^[60]^ 2g4l.^[61]^ The structure of salicylic acid binding protein 2 from tobacco without a bound ligand used for comparison has the protein data bank ID 1y7h.^[14]^ PyMOL v.2.5.4 was used to overlay the structures using the align function and to create images of protein structures.^[62]^

## Acknowledgements

Funding for this research was provided by US National Science Foundation award CBET-2039039, National Institutes of Health/National Institute of General Medical Sciences (grant No. GM119483); NIH (grant No. NIGMS R35-GM118047); X-ray diffraction data were collected at the Northeastern Collaborative Access Team beamlines, which are funded by the U.S. National Institutes of Health (NIGMS P30 GM124165). The Pilatus 6M detector on the 24-ID-C beamline is funded by a NIH-ORIP HEI grant (S10 RR029205). This research used resources of the Advanced Photon Source, a U.S. Department of Energy (DOE) Office of Science User Facility operated for the DOE Office of Science by Argonne National Laboratory under Contract No. DE-AC02-06CH11357. G.C. and S.O. thank the Generalitat de Catalunya for the consolidated group TCBioSys (SGR 2021 00487) and grant projects PID2021-129034NB-I00 and PDC2022-133950-I00 funded by Spanish MICIN. S.O. is grateful to the funding from the European Research Council (ERC) under the European Union’s Horizon 2020 research and innovation program (ERC-2015-StG-679001, ERC-2022-POC-101112805, and ERC-2022-CoG-101088032), and the Human Frontier Science Program (HFSP) for project grant RGP0054/2020. G. C. was supported by a research grant from ERC-StG (ERC-2015-StG-679001) and HFSP RGP0054/2020. We thank Drenen Magee for help with the analysis of x-ray crystallography data.

## Notes

https://www.rcsb.org/structure/8EUO

### Competing Interest Statement

The authors have declared no competing interest.

