## Supplementary Information for "Designing Efficient Enzymes: Eight Predicted Mutations Convert a Hydroxynitrile Lyase into an Efficient Esterase"

Supplementary Figures……s2
Supplementary Tables……s5
Supplementary References……s13

### Supplementary Figures

```
SABP2           1 MKEGKHFVLVHGACHGGWSWYKLKPLLEAAGHKVTALDLAASGTDLRKIE     50
                   ....||||:|..|||.|.|:||||||||.|||||||||||||.|.|:||
HbHNL           1 -MAFAHFVLIHTICHGAWIWHKLKPLLEALGHKVTALDLAASGVDPRQIE     49

SABP2          51 ELRTLYDYTLPLMELMESLSADEKVILVGHSLGGMNLGLAMEKYPQKIYA    100
                  |:.:..:|:.||:..:|:|...|||||||.|.||:|:.:|.:||.:||.|
HbHNL          50 EIGSFDEYSEPLLTFLEALPPGEKVILVGESCGGLNIAIAADKYCEKIAA     99

SABP2         101 AVFLAAFMPDSVHNSSFVLEQYNERTPAENWLDTQFLPYGSPEEPLTSMF    150
                  |||..:.:||:.|..|:|:::..|..|  :|.||.:..|....:.:|.:.
HbHNL         100 AVFHNSVLPDTEHCPSYVVDKLMEVFP--DWKDTTYFTYTKDGKEITGLK    147

SABP2         151 FGPKFLAHKLYQLCSPEDLALASSLVRPSSLFMEDLSKAKYFTDERFGSV    200
                  .|...|...||.||.||:..||..|.|..|||...|:|..:||.|.:||:
HbHNL         148 LGFTLLRENLYTLCGPEEYELAKMLTRKGSLFQNILAKRPFFTKEGYGSI    197

SABP2         201 KRVYIVCTEDKGIPEEFQRWQIDNIGVTEAIEIKGADHMAMLCEPQKLCA    250
                  |::|:...:|:....|||.|||:|....:..:::|.||...|.:.:::..
HbHNL         198 KKIYVWTDQDEIFLPEFQLWQIENYKPDKVYKVEGGDHKLQLTKTKEIAE    247

SABP2         251 SLLEIAHKYN   260
                  .|.|:|..||
HbHNL         248 ILQEVADTYN   257
```

**Supplementary Fig. 1 | Pairwise amino acid sequence alignment of SABP2 (UniProt Q6RYA0) and *Hb*HNL (UniProt P52704) using the Needleman-Wunsch algorithm.**[1] The comparison over 260 positions identified 114 (44%) identical positions (marked by '|'), 161 (62%) similar positions (identical plus those marked by ':') and 3 (1.2%) gaps (marked by a blank and a '-' in the *Hb*HNL sequence). The default settings (EBLOSUM62 matrix, gap penalty of 10, extend penalty of 0.5) of the web tool EMBOSS Needle (https://www.ebi.ac.uk/Tools/psa/emboss_needle/) were used to create the alignment.

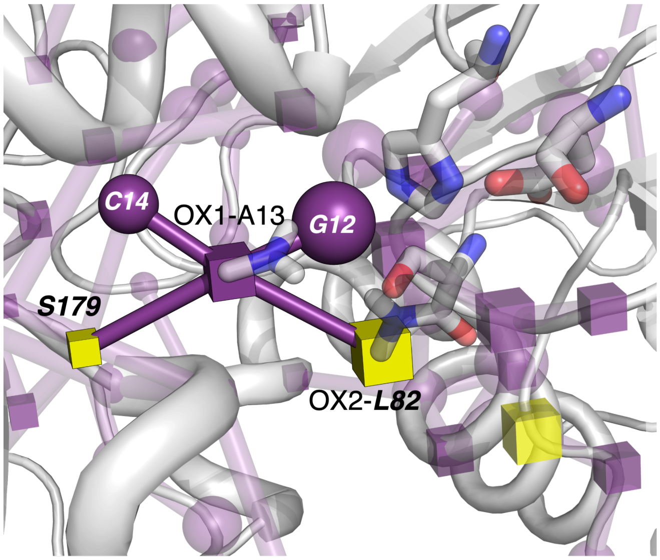

**Supplementary Fig. 2** | Zoom and slight rotation relative to Fig. 3 of the SPM of SABP2 showing the correlated motions of OX1 (Ala13) and residues Cys14, Gly12, Ser179, and OX2 (Leu 82). Cys14 and Gly12 are conserved between SABP2 and HNL3V (and are shown as spheres), whereas Ser179 and Leu82 (shown as cubes) correspond to Gly176 and Cys81 in HNL3V. The catalytic residues and amides of the oxyanion hole residues are shown in sticks. C14 and G12 were hard to see in Fig. 3.

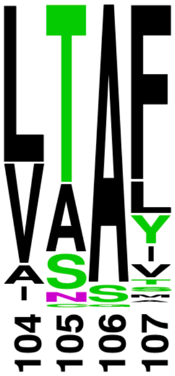

**Supplementary Fig. 3 | Relative frequencies of amino acids at positions 104-107 among 889 homologs of SABP2.** The height of the letters indicates the relative frequency of the amino acid. The positions correspond to positions 103-106 in *Hb*HNL. The most common amino acid at SABP2 position 105 is threonine; the second most common is alanine. The multiple sequence alignment was generated using Consensus Finder[2] (http://kazlab.umn.edu/) using default settings and the UniProt SABP2 sequence Q6RYA0. Consensus Finder searches for homologs using BLAST, reduces sequence redundancy by clustering all sequences into groups with 90% sequence identity using CD-HIT and retains only one representative sequence from each cluster. Finally, Clustal W creates the multiple sequence alignment. The image was generated with WebLogo[3] (https://weblogo.berkeley.edu/).

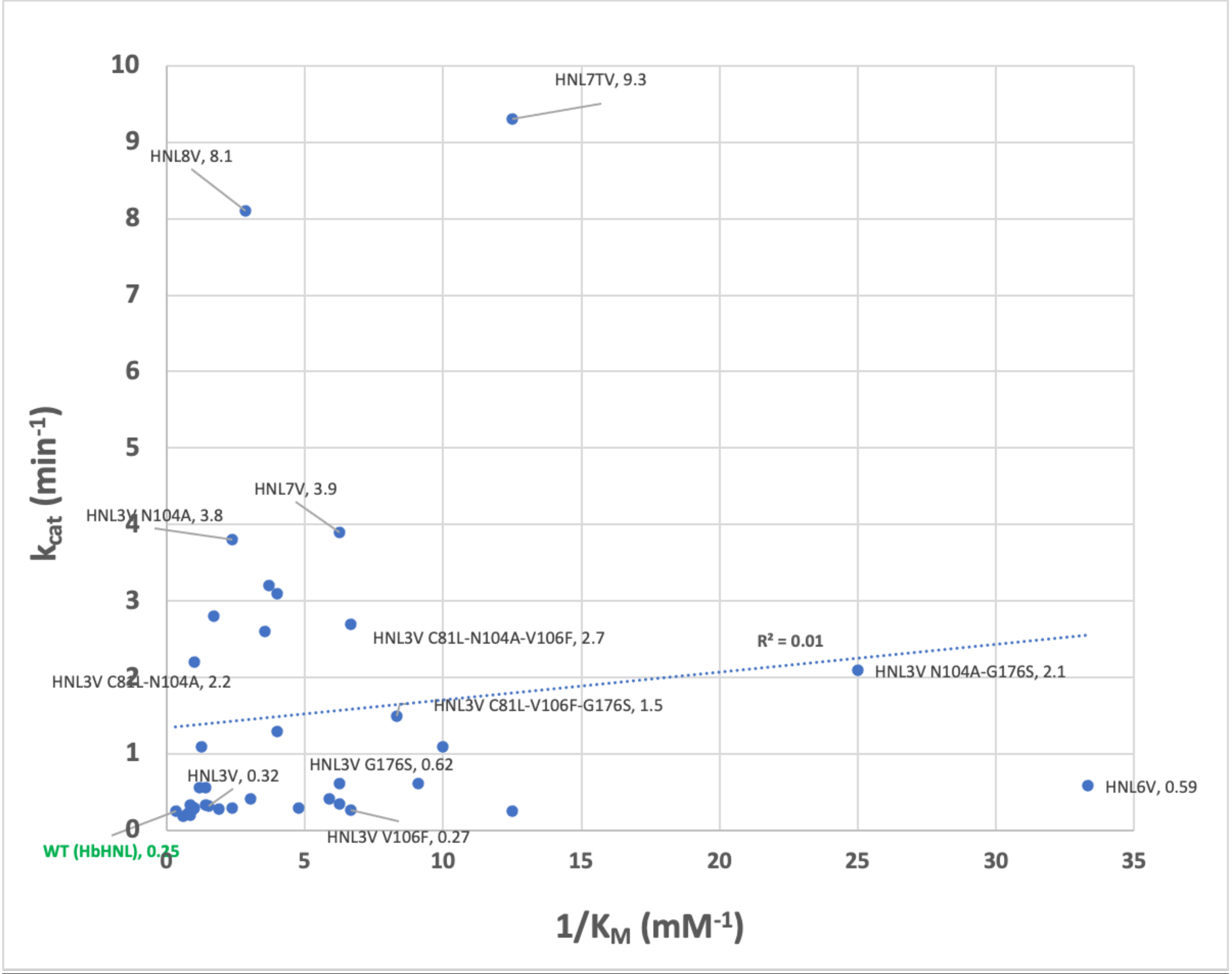

**Supplementary Fig. 4 | Improvements in catalytic turnover and binding are independent.** Linear regression of $k_{cat}$ (y-axis) vs. $1/K_M$ (x-axis) values for *Hb*HNL variants shows no correlation ($R^2 = 0.01$). $\mathbf{k_{cat}}$ values are shown for selected variants.

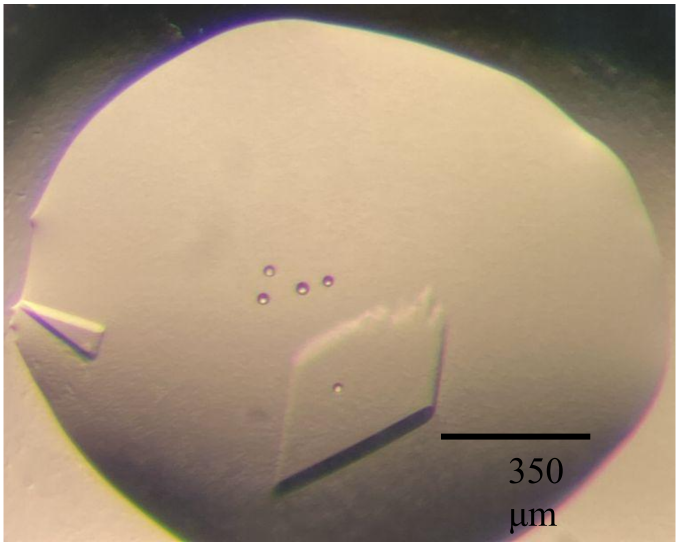

**Supplementary Fig. 5 | HNL6V crystal used for data collection prior to extraction or soaking.** Growth conditions: 0.1 *M* BIS TRIS, pH 5.5, 2.0 *M* ammonium sulfate. The HNL6V crystals measured 90x175 µm (left crystal) and 350x300 µm (right crystal). The right crystal was harvested for data collection. The five circular "bubbles" are optical artifacts of the microscope.

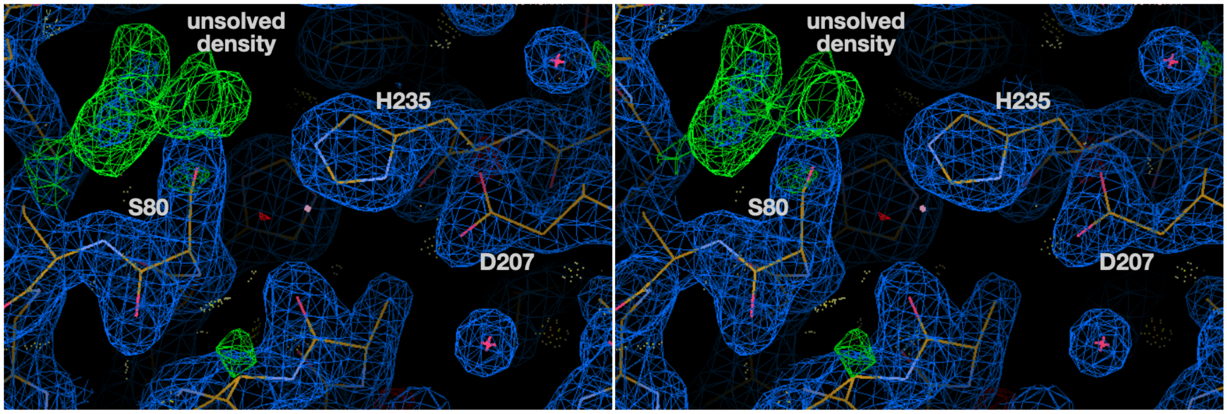

**Supplementary Fig. 6 | Cross-eyed stereo view of the unmodeled density (green mesh) of HNL6V near the active site at a 3.0 Å contour level.** The electron density associated with the catalytic triad (S80-D207-H235) is labeled.

### Supplementary Tables

**Supplementary Table 1 | Steady-state kinetic parameters for hydrolysis of *p*-nitrophenyl acetate of all enzyme variants.** See Materials and Methods for experimental conditions and details.

| Enzyme | $k_{cat}$ (min $^{-1}$) | $K_M$ (mM) | $k_{cat}/K_M$ ($M^{-1}$*$min^{-1}$) |
|---|---|---|---|
| SABP2 | 130±3.7 | 2.2±0.17 | 61,000 |
| WT (HbHNL) | 0.25±0.02 | 3±0.4 | 84 |
| HNL3 = WT T11G-E79H-K235M | 0.33±0.02 | 0.71±0.1 | 460 |
| HNL3V = HNL3 H103V | 0.32±0.02 | 0.65±0.1 | 490 |
| HNL3V C81A | 0.29±0.04 | 0.42±0.3 | 690 |
| HNL3V N104A | 3.8±0.23 | 0.42±0.08 | 9,100 |
| HNL3V G176S | 0.62±0.02 | 0.16±0.03 | 3,900 |
| HNL3V I12A | 0.28±0.03 | 0.53±0.17 | 530 |
| HNL3V C81L | 0.28±0.02 | 1±0.18 | 270 |
| HNL3V F54L | 0.35±0.01 | 0.16±0.03 | 2,200 |
| HNL3V V106F | 0.27±0.01 | 0.15±0.05 | 1,800 |
| HNL3V I209G | 0.61±0.07 | 0.11±0.08 | 5,500 |
| HNL3V I209G-F210I | 0.23±0.04 | 1.3±0.56 | 180 |
| HNL3V N104A-G176S | 2.1±0.06 | 0.04±0.01 | 52,000 |
| HNL3V C81L-N104A | 2.2±0.08 | 0.99±0.13 | 2,200 |

| **Enzyme** | **$k_{cat}$ (min $^{-1}$)** | **$K_M$ (mM)** | **$k_{cat}/K_M$ ($M^{-1}$*$min^{-1}$)** |
|---|---|---|---|
| HNL3V N104A-V106F | 3.2±0.22 | 0.27±0.08 | 12,000 |
| HNL3V V106F-G176S | 3.1±0.21 | 0.25±0.06 | 13,000 |
| HNL3V C81L-V106F | 1.3±0.03 | 0.25±0.03 | 5,300 |
| HNL3V C81A-G176S | 0.42±0.02 | 0.17±0.04 | 2,500 |
| HNL3V C81L-F54L | 0.41±0.02 | 0.33±0.06 | 1,200 |
| HNL3V C81L-F54L-V106F | 1.1±0.05 | 0.8±0.13 | 1,400 |
| HNL3V C81L-G176S | 0.3±0.02 | 1±0.23 | 300 |
| HNL3V C81L-F54L-G176S | 0.25±0.01 | 0.08±0.01 | 3,100 |
| HNL3V C81L-N104A-V106F | 2.7±0.18 | 0.15±0.05 | 18,000 |
| HNL3V N104A-V106F-G176S | 2.6±0.21 | 0.28±0.09 | 9,400 |
| HNL3V C81L-V106F-G176S | 1.5±0.02 | 0.12±0.01 | 13,000 |
| HNL3V C81L-I12A-V106F | 0.56±0.08 | 0.84±0.41 | 670 |
| HNL3V C81L-I12A-G176S | 0.3±0.01 | 0.21±0.05 | 1,400 |
| HNL3V C81L-F54L-V106F-G176S | 1.1±0.02 | 0.1±0.01 | 11,000 |
| HNL3V C81L-I12A-G176S-V106F | 0.56±0.06 | 0.71±0.26 | 790 |
| HNL3V C81L-G176S-V106F-I209G-F210I | 0.34±0.03 | 1.2±0.32 | 290 |
| HNL3V C81L-G176S-V106F-I209G-F210I-L121Y-F125T | 0.2±0.01 | 1.2±0.16 | 170 |

| Enzyme | $k_{cat}$ (min $^{-1}$) | $K_M$ (mM) | $k_{cat}/K_M$ ($M^{-1}$*$min^{-1}$) |
|---|---|---|---|
| HNL3V I209G-F210I-L121Y-F125T | 0.7 ± 0.11 | 0.97 ± 0.12 | 720 |
| HNL6V (HNL3V C81L-N104A-G176S) | 2.3±0.02[a] | 0.13±0.01[a] | 18,000[a] |
| HNL7V (HNL3V C81L-N104A-V106F-G176S) | 3.9±0.45 | 0.16±0.07 | 25,000 |
| HNL7TV (HNL3V C81L-N104T-V106F-G176S) | 9.3±0.33 | 0.08±0.04 | 120,000 |
| HNL8V (HNL3V C81L-N104A-S105A-V106F-G176S) | 8.1±0.38 | 0.35±0.06 | 23,000 |
| SABP2 A104N | 29±1.6 | 3±0.16 | 9,600 |

[a] Activity was measured at a higher temperature (29°C) relative to other variants (22 ±2 °C). Increased temperature correlates with higher observed reaction rates.

**Supplementary Table 2 | Rank of *Hb*HNL variants ordered by $k_{cat}$ and $K_M$**

| Ordered by $k_{cat}$ | | | Ordered by $K_M$ | |
|---|---|---|---|---|
| **Variant** | **$k_{cat}$ (min$^{-1}$)** | **Rank order** | **$K_M$ (mM)** | **Variant** |
| HNL7TV | 9.34 | 1 | 0.04 | HNL3V N104A-G176S |
| HNL8V | 8.13 | 2 | 0.08 | HNL7TV |
| HNL7V | 3.92 | 3 | 0.08 | HNL3V C81L-F54L-G176S |
| HNL3V N104A | 3.81 | 4 | 0.1 | HNL3V C81L-F54L-V106F-G176S |
| HNL3V N104A-V106F | 3.21 | 5 | 0.11 | HNL3V I209G |
| HNL3V V106F-G176S | 3.14 | 6 | 0.12 | HNL3V C81L-V106F-G176S |
| HNL3V C81L-N104A-V106F | 2.72 | 7 | 0.13 | HNL3V C81L-N104A-G176S (aka HNL6V) |

| **Ordered by $k_{cat}$** | | | **Ordered by $K_M$** | |
|---|---|---|---|---|
| **Variant** | **$k_{cat}$ (min$^{-1}$)** | **Rank order** | **$K_M$ (mM)** | **Variant** |
| HNL3V N104A-V106F-G176S | 2.63 | 8 | 0.15 | HNL3V C81L-N104A-V106F |
| HNL6V (HNL3V C81L-N104A-G176S) | 2.3 | 9 | 0.15 | HNL3V V106F |
| HNL3V C81L-N104A | 2.2 | 10 | 0.16 | HNL7V |
| HNL3V N104A-G176S | 2.08 | 11 | 0.16 | HNL3V G176S |
| HNL3V C81L-V106F-G176S | 1.52 | 12 | 0.16 | HNL3V F54L |
| HNL3V C81L-V106F | 1.33 | 13 | 0.17 | HNL3V C81A-G176S |
| HNL3V C81L-F54L-V106F | 1.12 | 14 | 0.21 | HNL3V C81L-I12A-G176S |
| HNL3V C81L-F54L-V106F-G176S | 1.08 | 15 | 0.25 | HNL3V V106F-G176S |
| HNL3V G176S | 0.62 | 16 | 0.25 | HNL3V C81L-V106F |
| HNL3V I209G | 0.61 | 17 | 0.27 | HNL3V N104A-V106F |
| HNL3V C81L-I12A-V106F | 0.56 | 18 | 0.28 | HNL3V N104A-V106F-G176S |
| HNL3V C81L-I12A-G176S-V106F | 0.56 | 19 | 0.33 | HNL3V C81L-F54L |
| HNL3V C81A-G176S | 0.42 | 20 | 0.35 | HNL8V |
| HNL3V C81L-F54L | 0.41 | 21 | 0.42 | HNL3V N104A |
| HNL3V F54L | 0.35 | 22 | 0.42 | HNL3V C81A |
| HNL3V C81L-G176S-V106F-I209G-F210I | 0.34 | 23 | 0.53 | HNL3V I12A |

| Ordered by $k_{cat}$ | | | Ordered by $K_M$ | |
|---|---|---|---|---|
| **Variant** | **$k_{cat}$ ($min^{-1}$)** | **Rank order** | **$K_M$ (mM)** | **Variant** |
| HNL3 | 0.33 | 24 | 0.65 | HNL3V |
| HNL3V | 0.32 | 25 | 0.71 | HNL3V C81L-I12A-G176S-V106F |
| HNL3V C81L-G176S | 0.3 | 26 | 0.71 | HNL3 |
| HNL3V C81L-I12A-G176S | 0.3 | 27 | 0.8 | HNL3V C81L-F54L-V106F |
| HNL3V C81A | 0.29 | 28 | 0.84 | HNL3V C81L-I12A-V106F |
| HNL3V I12A | 0.28 | 29 | 0.99 | HNL3V C81L-N104A |
| HNL3V C81L | 0.28 | 30 | 1 | HNL3V C81L-G176S |
| HNL3V V106F | 0.27 | 31 | 1.04 | HNL3V C81L |
| HNL3V C81L-F54L-G176S | 0.25 | 32 | 1.16 | HNL3V C81L-G176S-V106F-I209G-F210I-L121Y-F125T |
| HNL3V I209G-F210I | 0.23 | 33 | 1.17 | HNL3V C81L-G176S-V106F-I209G-F210I |
| HNL3V C81L-G176S-V106F-I209G-F210I-L121Y-F125T | 0.2 | 34 | 1.3 | HNL3V I209G-F210I |
| HNL3V I209G-F210I-L121Y-F125T | 0.19 | 35 | 1.7 | HNL3V I209G-F210I-L121Y-F125T |

**Supplementary Table 3 | Mutagenic primers for site-directed mutagenesis**

| Primer/sequence name | Primer sequence |
|---|---|
| I208G-F209I Rev | tcggtccacacataaattttc |
| I208G-F209I Fwd | ccaagacgaaggtattttacctgaatttcaactctgg |

| | |
|---|---|
| I208G Fwd | ccaagacgaaggtttttacctgaatttcaac |
| F209I Fwd | ccaagacgaaataattttacctgaatttc |
| C81L-L54F-G176S-H103V Fwd | gattggctcatttgatgagtattc |
| C81L-L54F-G176S-H103V Rev | tcctcaatttgccttggg |
| 81L-103V-104A-106F-176S Fwd | tgttttcgtcgcgtcatttttgccagacac |
| 81L-103V-104A-106F-176S Rev | gcagctgcaatcttttcac |
| 103V-104A Fwd | tgttttcgtcgcgtcagtattgccagacac |
| 103V-104A Rev | gcagctgcaatcttttcac |
| 103V-81L-104A-106F Fwd | tgttttcgtcgcgtcattcttgccagacac |
| 103V-81L-104A-106F Rev | gcagctgcaatcttttcac |
| 103V-C81L-N104T-V106F-G176S Fwd | tgttttcgtcacctcattcttgccag |
| 103V-C81L-N104T-V106F-G176S Rev | gcagctgcaatcttttcac |
| H103V_C81L_N104A_Fwd | tttcgtcgcgtcagtattgccagacacc |
| H103V_C81L_N104A_Rev | acagcagctgcaatcttttcacagtatttatcagc |
| 103V_N104A_V106F_Fwd | gtcattcttgccagacaccgagcac |
| 103V_N104A_V106F_Rev | gcgacgaaaacagcagctgcaatcttttc |
| C81L Fwd | cagcctgggaggactcaatatagcaattg |
| C81L Rev | tggccaaccagaatcaccttttcccc |

| | |
|---|---|
| SABP2 A104N Fwd | tgttttcttgaacgctttcatgcctg |
| SABP2 A104N Rev | gcagcatagatcttttgtg |
| SABP2 A104N Gibson vector Fwd | ttaaaggtgctgatcacatggcaatgctatg |
| SABP2 A104N Gibson vector Fwd | taaccttctcatctgctgaaagagattcc |
| SABP2 A104N Gibson gene Fwd | ttcagcagatgagaaggttatattagtggg |
| SABP2 A104N Gibson gene Rev | catgtgatcagcacctttaatctctattgc |
| C81A Fwd | attctggttggccatagcgctggaggactcaatatagc |
| C81L Fwd | tggccacagcctgggaggactcaatatag |
| C81L Rev | accagaatcaccttttcc |
| H103V-N104A-S105A-V106F Fwd | tgccagacaccgagcactgcccat |
| H103V-N104A-S105A-V106F Rev | agaatgccgcgacgaaaacagcagc |
| F54L Fwd | gattggctcactggatgagtattc |
| F54L Rev | tcctcaatttgccttggg |
| G176S Fwd | gacaaggaagagctcattatttcaaaatattttagc |
| G176S Rev | aacatcttcgccagttcatattc |
| H103V Fwd | gattgcagctgctgttttcgtcaattcagtattgccagac |
| H103V Rev | gtctggcaatactgaattgacgaaaacagcagctgcaatc |
| I12A Fwd | tattcatggcgcgtgccacggtgc |
| I12A Rev | agaacaaaatgagcgaatg |

| | |
|---|---|
| L121Y F125T Fwd | tatatggaggtgacccccgactggaaagacacc |
| L121Y F125T Rev | cttatccacgacgtaagatgggc |
| V106F Fwd | ccacaattcatttttgccagacac |
| V106F Rev | aaaacagcagctgcaatc |

**Supplementary Table 4 | Crystallization conditions for the x-ray structure determination of HNL6V**

| | |
|---|---|
| Method | Vapor diffusion, sitting drop |
| Plate type | CrystalMation Intelli-Plate 96-3 low-profile |
| Temperature (K) | 293 |
| Protein concentration (mg $ml^{-1}$) | 9.3 |
| Buffer composition of protein solution | 5 *mM* BES, pH 7.2 |
| Composition of reservoir solution | 0.1 *M* Bis-Tris, pH5.5, 2 *M* $(NH_4)_2SO_4$ |
| Volume and ratio of drop (nl) | 200, 1:1 (protein:screen solution) |
| Volume of reservoir (µl) | 50 |

**Supplementary Table 5 | Data collection and processing for the x-ray structure determination of HNL6V.** Values in parentheses are for the outer shell.

| | |
|---|---|
| X-ray source | APS BEAMLINE 24-ID-C |
| Wavelength (Å) | 0.979 Å |
| Detector | DECTRIS EIGER2 S 16M |
| Exposure Time (s) | 0.2 |
| Crystal-to-detector distance (cm) | 230 |
| Angle increment (°) | 0.2 |
| Resolution Range (Å) | 43.03 -1.99 (2.04-1.99) |
| Space Group | $C222_1$ |
| a, b, c (Å) | 47.054, 106.378, 128.396 |
| α, β, γ (°) | 90, 90, 90 |
| Matthews coefficient ($Å^3$Da-1) | 2.74 |
| Solvent Content (%) | 55.07 |

| | |
|---|---|
| Total reflections | 102923 (7877) |
| Unique Reflections | 20181 (1606) |
| Multiplicity | 5.1 |
| Mosaicity (°) | 0.2 |
| Completeness (%) | 89.43 (89.95) |
| (I/$\sigma$(I)) | 13.9 (2.3) |
| Wilson B Factor (Å$^2$) | 28.09 |
| Rmerge | 0.063 (0.703) |
| Rmeas | 0.077 (0.861) |
| Rp.i.m | 0.032 (0.490) |
| $CC_{1/2}$ | 0.998 (0.689) |

**Supplementary Table 6 | Structure refinement for the x-ray structure determination of HNL6V.** Values in parentheses are for the outer shell.

| | |
|---|---|
| Reflections used in refinement | 20178 (2005) |
| Reflections used for $R_{free}$ | 2000 (199) |
| $R_{work}$ | 0.1849 (0.2285) |
| $R_{free}$ | 0.2374 (0.2603) |
| No. of non-H atoms | |
| total | 2162 |
| Macromolecules | 2030 |
| Ligands | 0 |
| Solvent | 123 |
| No. of protein residues | 255 |
| R.m.s.d, bonds (Å) | 0.009 |
| R.m.s.d, angles (°) | 1.03 |
| Ramachandran favored (%) | 96.44 |
| Ramachandran preferred (%) | 96.44 |
| Ramachandran allowed (%) | 3.56 |

| | |
|---|---|
| Ramachandran outliers (%) | 0.00 |
